# Experimental platform for the functional investigation of membrane proteins in giant unilamellar vesicles

**DOI:** 10.1101/2021.12.22.473796

**Authors:** Nicolas Dolder, Philipp Müller, Christoph von Ballmoos

**Author notes:** Contributed equally to the manuscript (Nicolas Dolder, Philipp Müller).

## Abstract

Giant unilamellar vesicles (GUVs) are micrometer-sized model membrane systems that can be viewed directly under the microscope. They serve as scaffolds for the bottom-up creation of synthetic cells, targeted drug delivery and have been used in many *in vitro* studies of membrane related phenomena. GUVs are also of interest for the functional investigation of membrane proteins that carry out many key cellular functions. A major hurdle to a wider application of GUVs in this field is the diversity of existing protocols that are optimized for individual proteins. Here, we compare PVA assisted and electroformation techniques for GUV formation under physiologically relevant conditions, and analyze the effect of immobilization on vesicle structure and membrane tightness towards small substrates and protons. There, differences in terms of yield, size, and leakage of GUVs produced by PVA assisted swelling and electroformation were found, dependent on salt and buffer composition. Using fusion of oppositely charged membranes to reconstitute a model membrane protein, we find that empty vesicles and proteoliposomes show similar fusion behavior, which allows for a rapid estimation of protein incorporation using fluorescent lipids.

## Introduction

Giant unilamellar vesicles (GUVs) are micrometer sized (1-100 µm) model membrane systems serving for *in vitro* studies of various lipid membrane related processes, such as membrane fusion and fission, cell division or lipid domain formation.^1^ GUVs are also considered as scaffolds for the bottom-up creation of synthetic cells^2–4^ and the targeted delivery of drugs.^5^ Another, much less explored application of GUVs is the investigation of membrane proteins (MPs) that are responsible for many cellular key functions, such as nutrient uptake, signal transduction, energy metabolism and regulation of the structure and dynamics of the membrane.^6^ In humans, altered expression or activity of MPs is related to many diseases, such as neurodegenerative disorders, diabetes and certain cancers.^7–10^ Being accessible on the cellular surface, MPs are considered excellent drug targets.^11^ A profound understanding of the enzyme mechanism and a robust platform to investigate mechanism and potential drug targets are key to the successful development of new treatment methods. With their size close to eukaryotic cells that reduces curvature stress compared to small vesicles, GUVs are an attractive platform for MP analysis. Since measurements are possible on a single vesicle level, much less protein is required, opening this field also to poorly expressing human proteins.

In our laboratory, we investigate the molecular mechanism of different membrane proteins, especially solute transporters, and members of the respiratory chain from bacteria and eukaryotes.^12–14^ As others, we were attracted by the unique possibility of GUVs to follow the function of our proteins of interest, often including the transport of substrates as small as protons, directly under the microscope. The first functional reconstitution of a membrane protein into GUVs has been described already twenty years ago, but until today only around a dozen publications demonstrate convincing light microscopic measurements with such transporters.^15–25^ Although these publications impressively show the potential of the methods, applications beyond proof-of principle are rare. To understand the difficulty of these experiments, we have dissected them into the substeps (I) formation, (II) immobilization and proton leakage, and (III) protein reconstitution.

I. Vesicles should be produced in a suitable size range and yield that a sufficient number of free-standing GUVs can be identified as unilamellar vesicles in the field of view of the microscope. Very large vesicles are further undesirable due to concerns with stability and tightness of the membrane. A diameter of 5 – 20 µm is considered to be well suited for our applications, allowing for a number >20 vesicles in the 40x-field of view and compatible with the size range of many eukaryotic cells. Formation should further take place under physiologically relevant conditions, including relatively high salt and buffer concentrations that stabilize MPs and are used to generate electric Nernst potentials. Generally, solvent free methods would be preferred as traces thereof in the membrane (e.g. within the hydrophobic part of the bilayer) affects membrane thickness and is likely to negatively affect MP function. Details on various GUV formation methods have been described elsewhere.^26–29^
II. Immobilization of GUVs on the microscopy slide is a pre-requisite for the long-term observation of single vesicles to follow the function of MPs inside a GUV. Immobilization has been achieved using various strategies, both physical^30–34^ and non-specific^33,35–37^ or specific chemical interactions.^35^ Due to its high specificity and affinity (K_D_ ∼10^−15^ M), the biotin-avidin interaction has become a popular immobilization strategy.^37,38^ However, it has been shown that strong vesicle adhesion forms a spherical cap on the surface^39^ that can lead to an increase in membrane tension and formation of pores, making GUVs prone to leakage, or ultimately even leading to rupture of the vesicles.^39–41^ Such leakage is detrimental for measuring metabolic transporters and especially respiratory enzymes, or if the membrane should be able to hold a membrane potential or proton gradient. In general, relatively little data is available regarding the tightness of the GUV membrane towards certain substrates,^42,43^ especially small ones such as protons.^25^
III. Functional reconstitution of the MP into the GUV membrane is critical for all downstream applications. Several methods including detergent-mediated reconstitution, rehydration of partially dehydrated SUVs or fusion of SUVs to GUVs have been exploited for MP reconstitution in GUVs.^44^ We and others have recently used charge-mediated fusion to introduce MPs into GUVs.^45–47^ This reconstitution strategy is a mild approach that transfers one or more MPs from SUVs containing positively charged lipids (reconstituted with traditional methods) to GUVs containing negatively charged lipids,^45,46^ preserving the protein orientation. This fusion strategy is not without concerns, and its limitation are discussed later.

Here, using polymer assisted swelling^48–51^ and electroformation^52^ methods, we compare the produced GUVs on vesicle properties important for investigating MPs in GUVs as outlined above. We compare the ability of these formation methods to produce a high number of vesicles with diameters between 5 – 20 µm under physiologically relevant conditions and characterize whether immobilization using the biotin-streptavidin system was affected by the presence of salt. Vesicle leakage is analyzed using influx measurements of the hydrophilic dye pyranine (HPTS). GUVs that did not show HPTS leakage were further analyzed using a proton efflux assay to assess the ability of GUVs to maintain a pH gradient. As a last step, we used fluorescently labeled cytochrome *bo*_*3*_ ubiquinol oxidase as a model protein to monitor incorporation via charge mediated fusion.

## Results

### GUV formation

The best-known method for GUV formation is based on electroformation, in which hydration of dried lipids is aided by an oscillating electrical field with different frequencies and voltages. More recently, polymer assisted swelling was reported as a simpler and versatile alternative and a variety of polymers have been used as substrate.^48–51,53,54^ Especially, production of GUVs with various lipid and buffer compositions was possible without the need for optimizing the method.^48–51^ This is in contrast to electroformation where protocols typically must be adapted, i.e. to form GUVs in buffers containing physiological salt concentrations^50,55–57^ or using cationic lipids.^58^ This makes polymer assisted swelling a desirable method for the investigation of MPs as variation of the lipid and buffer composition are common requirements to obtain optimal enzyme activity or to investigate lipid effects. Among the different substrates used, polyvinyl alcohol (PVA) has established itself as a go-to material due to its wide accessibility and ease of use. In the following, we have thus focused on the use of PVA and compared GUV properties important for MP investigation for vesicles produced by PVA assisted swelling and electroformation. As lipid composition, we chose 70% DOPC and 30 % DOPG, yielding a lipid bilayer that is negatively charged, comparable to bacterial or mitochondrial membranes. However, other lipid compositions should work analogously.

First, GUV formation was compared in the absence or presence of up to 100 mM monovalent salts (sodium and potassium chloride) and 10 mM divalent salt (calcium chloride), which we consider the upper limit of physiological concentrations needed for MP investigation (Figure 1). As most vesicles formed were unilamellar by eye, this property was ignored and only yield and size distribution of GUVs formed by the different methods were analyzed. Using 20 μg lipid as starting material, GUV formation by PVA assisted swelling and electroformation, either using platinum (Pt) wires or indium-tin-oxide (ITO) coated glass slides were compared (see Experimental procedures and Figure S1 for sequence protocols and chamber designs). As MPs are usually investigated in free-standing vesicles, yield and size distribution were characterized for carefully detached GUVs transferred from the formation chamber to 8 well chambered microscopy slides. Our focus was the number of GUVs with diameters between 5 and 20 µm that we considered suitable for our purposes (see above). GUV formations were considered as not successful if <10 GUVs in the desired size range were observed per field of view under the same experimental formation conditions. In the following, we considered 1.5× 10^5^ GUVs per ml in the requested size range as sufficient. The number is based on the observation that loading of 100 µL vesicle suspension should lead to at least 20 GUVs per field of view.

**Figure 1:**
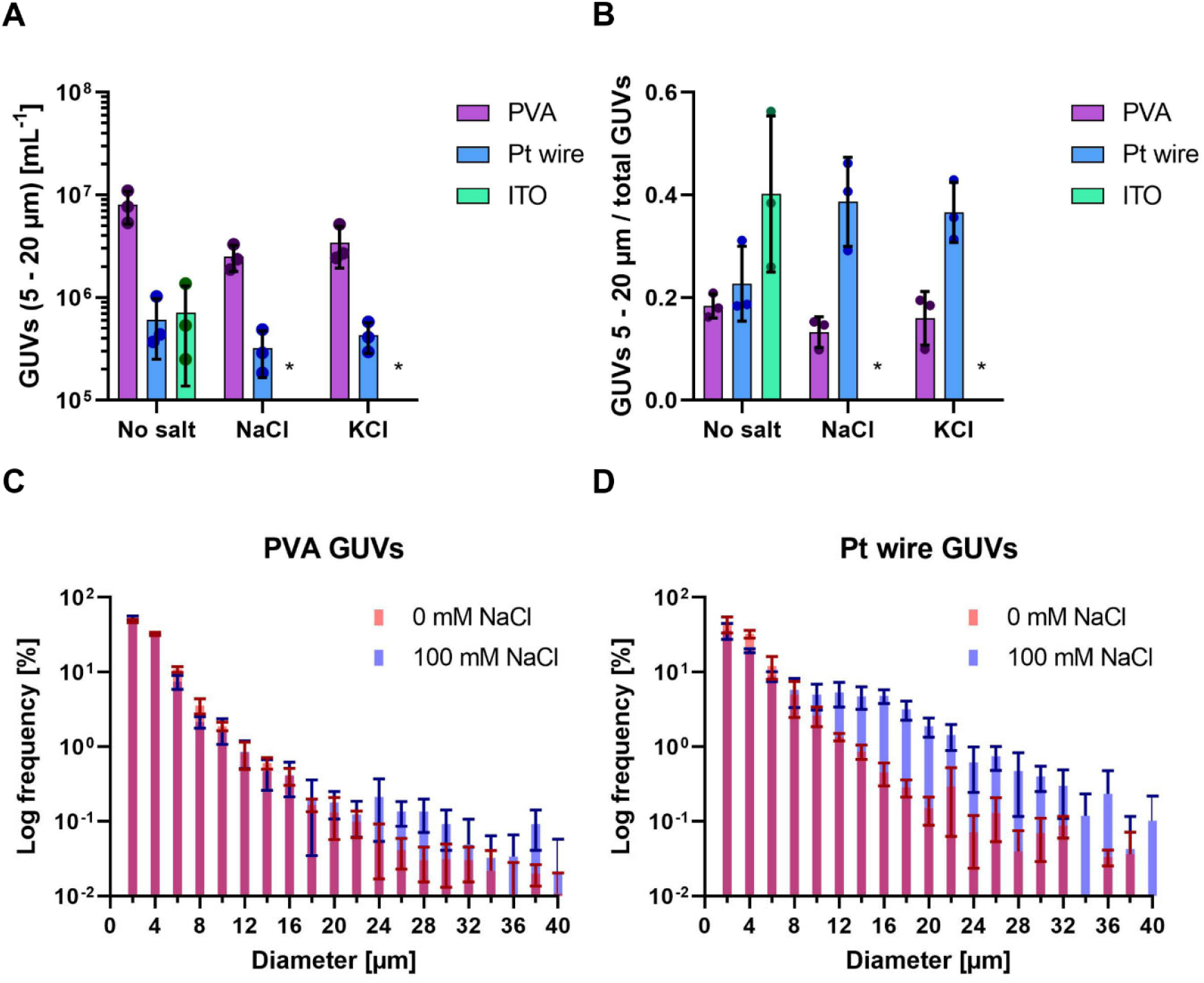
Characterization of GUV electroformation and PVA-assisted swelling. For each condition, size and concentration of GUVs was obtained from sampling of 9 positions in each well, conducted in three independent experiments and analyzed as previously published.^59^ GUV formations that produced an insufficient number of vesicles (< 10 GUVs in the requested size range per 40x field of view) are marked by an asterisk. The height of the bar indicates the average with individual values from the experiments shown as dots. Error bars indicate the standard deviation. A) Concentration of GUVs with diameters of 5 – 20 µm in solution after removal from the formation chamber. B) Formation quality assessed by calculating the fraction of GUVs in the desired diameter range compared to the total amount of GUVs counted. C-D) Histograms of PVA formation and Pt wire formation in the presence or absence of 100 mM NaCl. Diameters were obtained from 780 – 6480 GUVs depending on the formation. The width of the bins is 2 µm and the graph is cut off at 40 µm. Frequencies are displayed on a logarithmic scale as larger vesicles have smaller frequencies. C) Histograms of GUVs prepared by PVA formation. D) Histograms of GUVs prepared by Pt wire electroformation.

In the absence of salt, high concentrations (>7 × 10^5^/ml) of GUVs were obtained for all three tested formation methods. In the presence of 100 mM monovalent salts NaCl or KCl, however, only formation via PVA assisted swelling and Pt wire formation was satisfactory. Insufficient GUV concentrations were also obtained for 10 mM calcium chloride for all three formations (Figure S2). In all conditions, PVA assisted swelling yielded the largest number of GUVs in the desired size range. In the absence of salt, ∼8 × 10^6^ GUVs (5 – 20 µm) per 1 mL formation solution (Figure 1A) were obtained, while yields decreased 2 to 3-fold in presence of 100 mM monovalent salt. Roughly, the number of GUVs obtained from both electroformation methods at 0 mM salt was about 10 times smaller than that for PVA, yielding still sufficiently high concentrations for downstream applications. Interestingly, the impact of 100 mM salt addition was less pronounced using Pt wire formation compared to the PVA method. To assess how well each formation method is suited to obtain GUVs in the desired size range, the fraction of GUVs formed with diameters of 5 - 20 µm relative to the total number was calculated (Figure 1B). The smallest fractions (<20%) were obtained for all three PVA formations, indicating a high number of small vesicles, while large fractions (∼40%) were obtained for Pt wire formation in the presence of 100 mM monovalent salt and ITO formation in absence of salt. Finally, histograms (bin size = 2 μm) of the size distribution for PVA GUVs and Pt wire GUVs with and without NaCl were calculated (Figure 1C and D) as well as the lipid yield and average diameter (Table S1), which are presented in more detail in the supplementary information. In the absence of salt, PVA and Pt wire GUVs showed a similar size distribution, while a larger frequency of GUVs >10 µm was obtained for ITO formation (Figure S3). Interestingly, an increased frequency of larger GUVs (> 10 µm) is observed for Pt wire formation in the presence of NaCl compared to no salt, while PVA formation was barely affected, except for a slight increase for GUVs > 20 µm. Both NaCl and KCl behaved similarly in either PVA or Pt wire formation (Figure S4).

### GUV immobilization

Typically, spectroscopic assays for MPs reconstituted in SUVs last from a few seconds to at least several minutes. Often, at one or more timepoints, substrates or other chemicals are added to start the reaction or otherwise influence it. It is therefore indispensable to immobilize GUVs to observe them in the same field of view over a prolonged time. Here, we focus on the well-known interaction between biotin and streptavidin that is frequently used as an immobilization system, in which biotinylated lipids in the GUV membrane interact with the streptavidin coated surface of a microscopy slide. The system allows to fine-tune the degree of immobilization by varying the concentration of either biotinylated lipids or streptavidin. We anticipate that also the ionic strength of the surrounding buffer might influence the immobilization behavior of GUVs. To standardize the immobilization condition in different microscopy slides, we calculated the streptavidin amount per area of slide surface in contact with the streptavidin solution, and the value was termed “streptavidin density” in [ng mm^-2^]and is used from now on. In the next series of experiments, GUVs produced via the PVA and Pt wire method were immobilized using two different streptavidin densities, both in the presence and absence of 100 mM NaCl. As shown in Figure 2, a spherical adhesion cap was observed for both formation methods at high streptavidin density, and the presence of NaCl promoted the formation of adhesion caps at low streptavidin density. No caps were observed without streptavidin (Figure S5A) in either condition. Since cap sizes within one condition were rather heterogenous (Figure S5B), quantification of immobilization was not straightforward. Using commercial channel slides, we therefore developed a flow-based assay that greatly facilitated quantification of the immobilization behavior (Figure 3A). Since PVA and Pt wire GUVs showed comparable immobilization in 8 well chambered slides (Figure 2), only PVA GUVs were used in flow experiments due to the higher concentration after formation.

**Figure 2.**
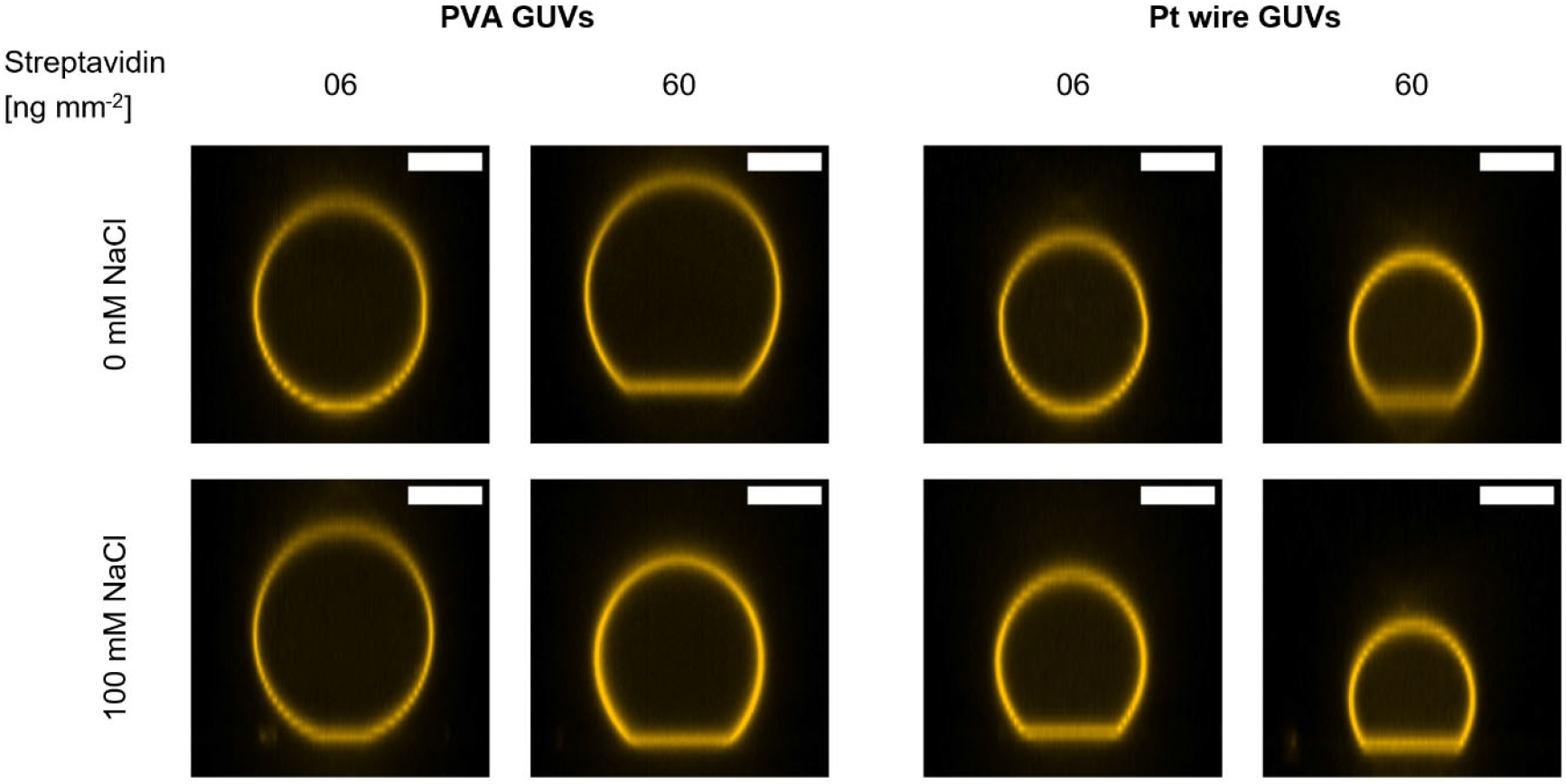
Formation of spherical adhesion cap at low and high streptavidin density. PVA and Pt wire GUVs in the presence and absence of 100 mM NaCl were immobilized at different streptavidin densities and confocal Z-stacks were recorded. Side-views of representative GUVs from each condition are shown. Spherical caps are observed for all conditions at high streptavidin densities while low densities show no or few caps in the absence of NaCl. The scale bar is 10 µm.

**Figure 3.**
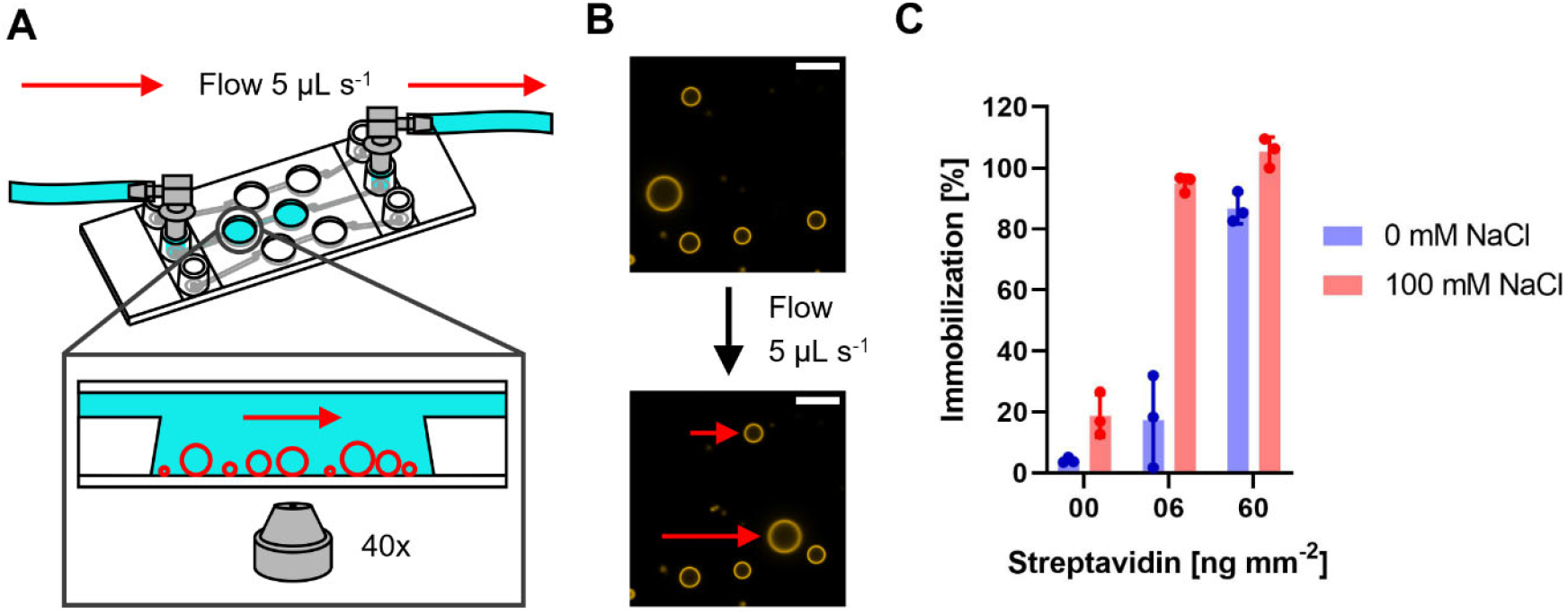
Immobilization assay using channel slides and PVA GUVs. A) Schematic representation of the channel slide setup with a blow-up depicting the immobilized GUVs (red circles) in the well of the channel. B) Example of immobilized GUVs under flow. Top image depicts GUVs before and bottom image during application of flow. Non-immobilized GUVs are indicated by the red arrows. The scale bar is 10 µm. C) Percentage of immobilized PVA GUVs in the presence or absence of 100 mM NaCl at different streptavidin densities assessed by comparing the number of immobilized GUVs and the number of GUVs before application of flow. Data from three independent experiments are shown. 40 – 120 GUVs were counted for analysis. The height of the bar indicates the average percentage of immobilization with individual values from the experiments shown as dots. Error bars indicate the standard deviation.

In this assay, PVA formed GUVs were immobilized at different streptavidin densities in presence and absence of 100 mM NaCl. First, all GUVs in the field of view were counted, before a constant flow of 5 µl s^-1^ was applied and the number of GUVs which withstood the flow were counted to determine the immobilization percentage. The general trend of the 8 well slide experiments was also observed in the flow-based immobilization assay. Without streptavidin, no immobilization was observed and at high streptavidin density almost all observed GUVs were fully immobilized in presence as well as in absence of NaCl (Figure 3C). However, GUVs immobilized at 6 ng mm^-2^ streptavidin in absence of NaCl showed almost no immobilization, whereas GUVs immobilized at the same streptavidin density but in presence of NaCl where immobilized almost to 100%. This indicates that immobilization using the biotin streptavidin system is highly salt sensitive.

### HPTS and proton leakage

To observe whether strong adhesion at high streptavidin densities affects the integrity of immobilized GUVs, we performed a dye permeation assay using HPTS. Empty GUVs were allowed to immobilize in the presence of HPTS on the outside and dye leakage into vesicles was monitored. At low streptavidin concentrations, almost no leakage was observed, while the fraction of leaky GUV increased with increasing streptavidin concentration, but never exceed 35% (Figure S6 and S8). In addition, slight differences were observed for PVA and Pt wire GUVs. For PVA GUVs, the optimal streptavidin density was found to be between 6 and 30 ng mm^-2^ and PVA GUVs in the presence of salt remained leaky after immobilization (Figure S7 and S8). The results are discussed in more detail in the Supplementary information. HPTS has a molecular weight of 524 g/mol, making it much larger than many biological substrates of interest. Protons, the smallest biological unit, are of special interest in our research with respiratory enzymes and we therefore extended our leakiness tests with GUVs to protons using a gramicidin assay (Figure 4). Here, GUVs containing the ratiometric proton sensitive fluorophore HPTS,^60–62^ which reports pH changes in the GUV lumen, are subjected to a proton gradient. Ratiometric dyes are attractive, as bleaching effects are suppressed and with HPTS, an increase of the pH in the GUV lumen leads to an increase in the ratio of the two signals. To limit the amount of buffering molecules, we used MOPS-KOH buffer instead of MOPS-BTP. Surprisingly, PVA GUVs prepared in 5 mM MOPS-KOH in the absence of salt showed insufficient immobilization even at high streptavidin densities (Figure S9), but immobilization was possible using 10 mM MOPS-KOH and 10 ng mm^-2^ streptavidin. To detect proton leakage, GUVs containing HPTS were prepared at pH 7.4, immobilized and washed with buffer at pH 8.0, leading to a gradient of 0.6 (inside acidic). The HPTS signal was monitored for 60 min before the pH gradient was equilibrated by the addition of the protonophore gramicidin, leading to an alkalinization of the GUV lumen and an increase in HPTS ratio (Figure S10). No change in signal was observed after gramicidin addition in the absence of a pH gradient (Figure 4A and C, Figure S10E and F).

Experiments with MPs are likely to last less than one hour. Thus, we analyzed only the increase in HPTS ratio in the first 20 min and compared them to the total increase after addition of gramicidin (Figure 4A). Few GUVs showed a decrease in HPTS ratio in the first 20 min and were excluded from the analysis. Decreasing signals might be observed in GUVs with HPTS leakage before or during incubation or due to microscope drift. With the remaining GUVs, the percentage of the increase after 20 min was calculated, assuming a total pH equilibration after gramicidin addition (Figure 4B). Most GUVs that were analyzed showed very little increase after 20 min. In those, proton leakage of less than 20% in the first 20 min was generally observed, indicating that the vesicles are sufficiently tight towards proton efflux if HPTS has not leaked. A slightly lower percentage of leaky GUVs was found in the PVA than in the Pt-wire preparation, and no influence of salt was observed in either of the preparations. A more detailed discussion of the data is found in the supplementary information.

### Membrane protein reconstitution

The overall negative surface charge of lipid composition of PC:PG (7:3) not only reflects conditions found in biological membranes, but is also compatible with fusion of oppositely charged proteo-SUVs.^45–47^ Here, every MP can be individually reconstituted into SUVs containing positively charged lipids under optimal conditions and possibly with a desired orientation. If these SUVs are mixed with negatively charged GUVs, membrane fusion occurs and the orientation of the membrane protein is conserved.^45^ This modular approach is ideally suited for the bottom-up construction of artificial cells that contain a variety of membrane proteins that have non-compatible reconstitution procedures. The reconstitution yield can be directly monitored, if the membrane protein is fluorescently labeled itself. However, this is not always feasible (impaired activity after labeling, low amount of protein, no suitable labeling chemistry). We therefore hypothesized that protein reconstitution can be correlated with fusion efficiency (which can be followed by fluorescent lipids). We were also interested to see if fusion of oppositely charged vesicles is dependent on the salt concentration, a topic that has been discussed with some controversy in the literature.^45,46,63^ To tackle the latter question, we thus investigated charge mediated fusion using protein free SUVs containing a lipid coupled dye with unlabeled GUVs in presence and absence of salt. With this approach, fusion was quantified by the amount of lipid coupled dye found in the previously unlabeled GUV membrane. If experiments were performed directly under the microscope using immobilized GUVs, rapid fusion is observed (Supplementary movie S1) with positively charged SUVs while no fusion was observed with neutral SUVs (Figure S11A). GUVs were fusogenic both in presence and absence of salt, although a smaller fluorescence increase in the GUV membrane was observed in the presence of salt (Figure 5A and S11B). However, the signal distribution in the raw data was rather heterogenous and we were not convinced of a significant influence of salt on the fusion process (Figure S12 and S13A). To eliminate effects of immobilization and mixing or pipetting artefacts during addition under the microscope, we repeated the experiments by mixing SUVs and non-immobilized GUVs in an Eppendorf tube (Figure S13B and S14). While PVA GUVs still show a slightly decreased fusion behavior in presence of salt, the same trend was not observed with PT wire GUVs (Figure S13B). The difference in observed fusion behavior of Pt-wire GUVs either immobilized on slides or in solution is not straightforward to explain. Additions to immobilized GUVs are performed with utmost care and mixing and diffusion of added chemicals are likely to differ from experiment to experiment. A detailed description is given in the supplementary information. Nevertheless, GUVs appeared fusogenic under all conditions. Next, we investigated fusion of proteoliposomes containing rhodamine-labeled lipids and DY-647P1-labelled *bo*_*3*_ oxidase in the absence of salt with PVA GUVs that showed a slightly higher fusion yield in the previous experiment (Figure 5B and C and Figure S15). Prior to fusion, proteoliposomes (protein and lipid labeled) and empty liposomes (only lipid labeled) were mixed in different ratios (3/0, 2/1, 1/2 and 0/3) to simulate the insertion of different amounts of enzymes. As depicted in Figure 5B, a linear correlation between *bo*_*3*_ oxidase signal and lipid-coupled dye was observed in the different experiments. This is a strong indication that empty SUVs and enzyme containing vesicles have the same fusion properties and that the intensity of lipid-coupled dye correlates with the intensity of labeled *bo*_*3*_ oxidase and can be used to estimate the relative amount of enzyme reconstituted.

**Figure 5:**
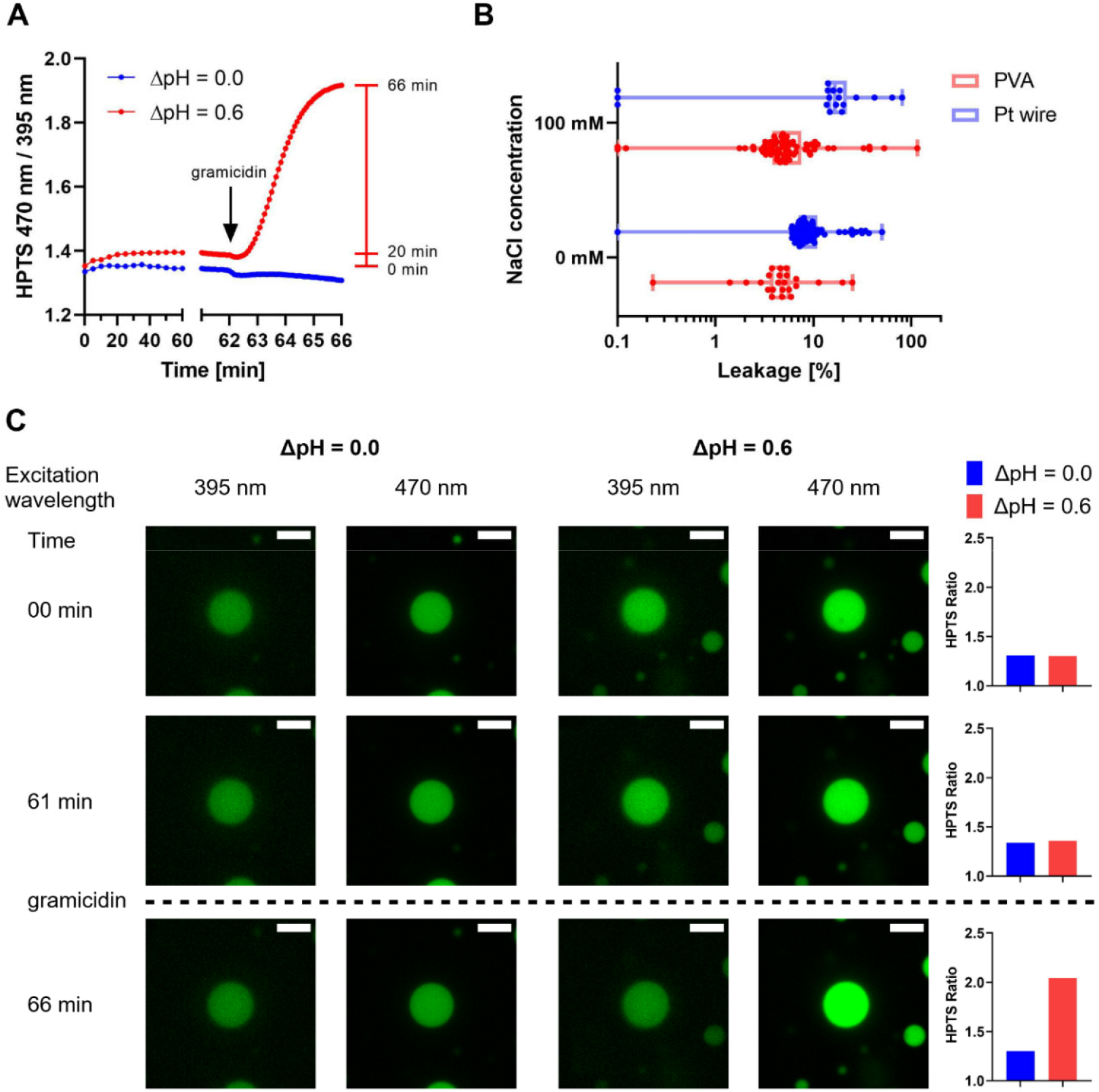
Proton leakage of immobilized PVA and Pt wire GUVs in the presence or absence of 100 mM NaCl. Analysis was limited to GUVs with diameters of 5 – 20 µm in the focal plane. Data are taken from a single time series for each condition. A) Average HPTS ratio of PVA GUVs with 100 mM NaCl subjected to a pH gradient of 0.0 and 0.6 over time (pH inside = 7.4, pH outside = 8.0). After 62 min, gramicidin was added to equilibrate the inner and outer pH, leading to an efflux of protons and an increase in HPTS ratio if a pH gradient is applied while no change is observed without pH gradient. The red bar indicates the time points used to assess the percentage of leakage. B) Box plot showing the percentage of leakage of PVA (red) and Pt wire GUVs (blue) with individual GUVs indicated as dots. Proton leakage was calculated by dividing the increase of the HPTS ratio after 20 min by the total increase from 0 min to 66 min (after addition of gramicidin). 20 – 70 GUVs per field of view were considered for analysis. C) Confocal microscopy images of an exemplary PVA GUV with 100 mM NaCl from the measurement with or without a pH gradient shown in (A). The two HPTS channels (green) are shown at different time points during the experiment. GUVs are shown at the start (00 min), after 1h incubation before (61 min) and after (66 min) the addition of gramicidin. On the right, the HTPS ratio of the GUVs depicted on the left were calculated. The scale bar is 10 µm. All images were processed identically.

**Figure 6:**
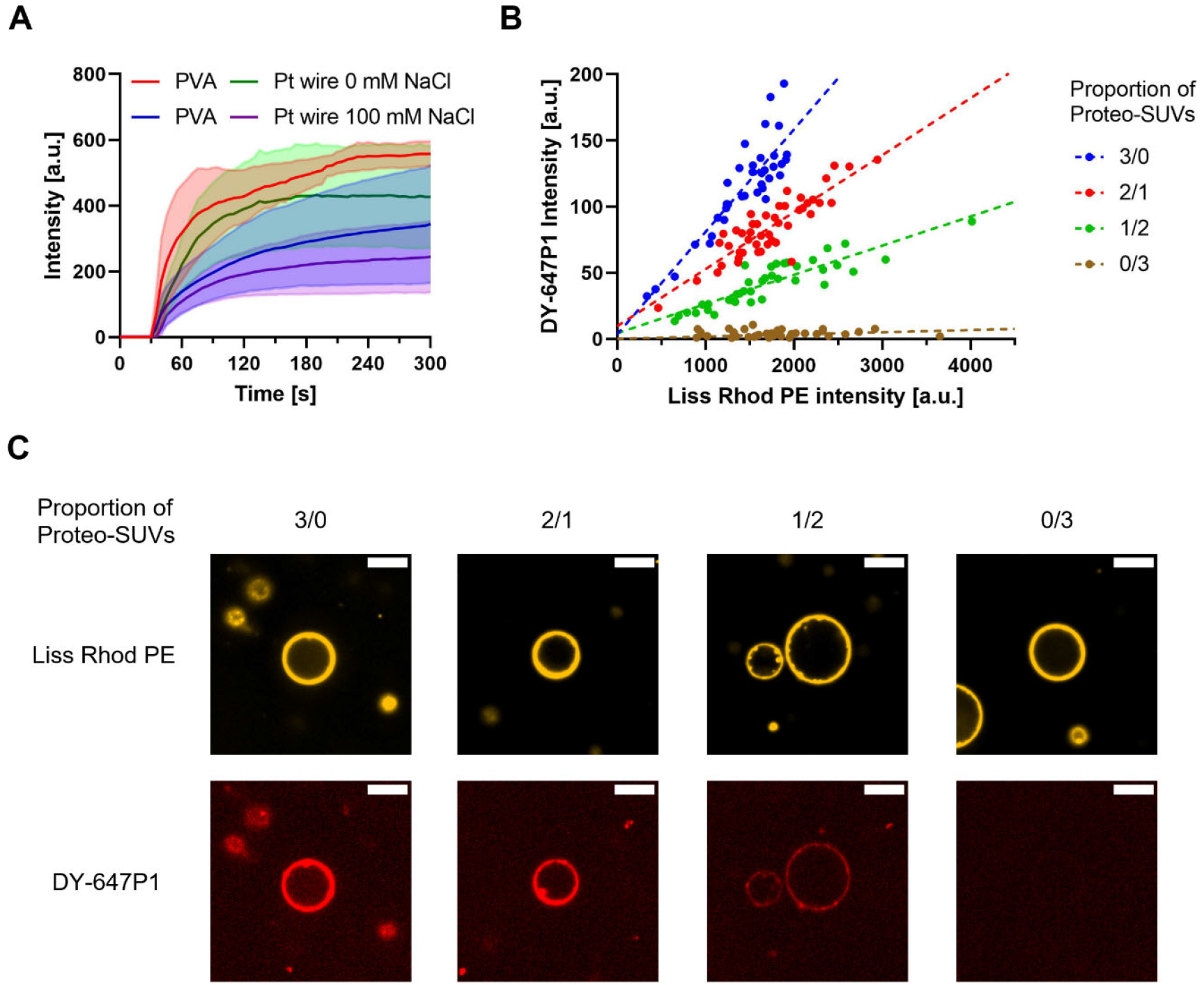
Charge mediated fusion of positively charged SUVs with negatively charged GUVs. Fusion was performed in an 8 well chambered slide with addition of SUVs to immobilized GUVs. Only GUVs with diameters of 5 – 20 µm in the focal plane were analyzed. A) Fusion of GUVs with empty SUVs (final concentration 10 µg mL^-1^) followed in real time. SUVs were added to the well after 30 s after which an increase of Liss Rhod PE signal in the GUV membrane was observed. Traces represent the mean (solid line) and standard deviation (transparent area above and below the trace) of the average intensity increase of 10 – 50 GUVs from 4 experiments. B) Fusion of PVA GUVs at 0 mM NaCl with empty SUVs and proteo-SUVs (final concentration 40 µg mL^-1^) containing DY-647P1-labeled cytochrome *bo*_*3*_ ubiquinol oxidase. Empty and proteo-SUVs were mixed at different ratios with proteo-SUV proportions of 3/0, 2/1, 1/2 and 0/3. The DY-647P1 intensity is compared to the Liss Rhod PE intensity 150 s after addition of SUVs. A similar distribution of Liss Rhod PE intensities is observed in all fusion reactions as both empty and proteo-SUVs contain Liss Rhod PE. The DY-647P1 shows a linear dependency to the Liss Rhod signal with a decreasing slope as the proportion of proteo-SUVs decreases. Values of individual GUVs from one experiment are shown (30 – 50 GUVs) as dots and the linear regression is represented as a dotted line. C) Confocal microscopy images of representative GUVs from fusion with different proteo-SUV proportions shown in (B). For each GUV, the Liss Rhod PE and DY-647P1 channel are depicted. The scale bar is 10 µm. All images from each channel were processed identically.

## Discussion

GUVs are attractive model systems for a variety of biological questions, including the investigation of membrane proteins. Except for a few examples, however, these investigations stayed at the level of proof-of-principle. In our lab, we are investigating different membrane proteins with substrates as small as proton or sodium ions, requiring robust and tight vesicles, also in the presence of physiological salt concentrations. As additions of substrates during the measurements are required, GUVs need to be immobilized to be monitored over a prolonged time. Reconstitution yield of membrane proteins into GUVs should be predictable to compare activities from single GUVs allowing to divide them into subpopulations. All these factors are highly important if GUVs should serve as a versatile and powerful model system to functionally characterize a MP beyond binding studies and serving as a spherical container.

Here, we compared the popular electroformation method with the more recently developed polymer assisted swelling for GUV formation in the presence of mono and -divalent ions. For electroformation, either Pt wires or ITO coated glass slides were used. All three methods yielded GUVs using buffer without salt with polymer assisted swelling producing the highest concentration and yield of GUVs. PVA assisted swelling and electroformation have been compared in the literature^50,59,64^ and the present data extend this knowledge with our comprehensive quantitative analysis using conditions for biological questions. In the presence of monovalent salt, GUVs were only obtained from PVA and Pt wire formation. This was not unexpected, as we used an electrode distance below 2 mm in our ITO coated glass slides that was previously shown to be insufficient for GUV formation at high salt concentrations.^57^ Overall, PVA assisted GUV formation in the presence of salt tended to decrease the concentration of vesicles, and is in agreement with other reports for polymer assisted swelling methods with increasing ionic strength or osmolarity of solution,^51,65^ although few polymer substrates showed increasing GUV concentrations.^66^ Thus, concentration of GUVs in the presence of salt might depend on the substrate which is used. Only slight decrease in vesicle concentration was observed in Pt-wire electroformation experiments, mostly due to an increase in vesicle diameter, as was previously observed for electroformation using up to 100 mM NaCl.^56^ We have used the same protocol as in the experiment without salt and the different sizes and concentration are likely a result of the changed electrical properties of the different formation conditions.^67^ No influence of salt on the size was observed for PVA assisted swelling, although a study using agarose had also observed an increase in GUV diameter with increasing ionic strength.^65^

The immobilization behavior of PVA and Pt wire GUVs was investigated using the widely used biotin streptavidin system in presence and absence of salt. It has been shown that strong adhesion of vesicles to a slide surface results in the formation of a spherical adhesion cap.^37,39^ Vesicle attachment to the slide surface is determined by the competition between the adhesive strength (that is the membrane-substrate interaction) and the bending rigidity of the vesicle membrane, the former one being sufficiently large or the latter one sufficiently small.^1^ During this process, leakage through water permeation or through the formation of pores might occur to adjust for the shape changes induced by strong adhesion.^39,40^ In agreement, strong adhesion has been shown to increase the membrane tension in GUVs^68,69^ with tension being a potential driving force for the formation of transient pores.^41^ Careful control of vesicle adhesion guarantees immobilization of GUVs while not compromising the membrane integrity. Modulation of the adhesion cap size was demonstrated by adjusting the MgCl_2_ concentration in the medium,^68^ which is also likely for other ions.^70^ Here, we found that the formation of a spherical adhesion cap greatly depends on salinity of the buffer as well as the amount of streptavidin used for immobilization. Immobilization was strictly dependent on streptavidin, as no unspecific binding was observed on BSA coated slides. Generally, the presence of salt increased the immobilization of vesicles and spherical cap size compared to the same conditions without salt. In our flow-based assay, we find that at the lowest tested streptavidin density of 6 ng mm^-2^, 20% and 100% of GUVs were immobilized in the absence or presence of NaCl, respectively. We also found an influence of the counterion (KOH or BTP) of the buffer (MOPS) if no salt was used, and that a higher buffer concentration improved immobilization. We consider the following two factors to be responsible for our observations. First, the presence of salt or other charged molecules in the buffer might influence the membrane-surface interaction, e.g. by screening repulsive charges or by bridging charged species on the surface and in the lipid membrane. It has been shown that salt facilitates the formation of supported lipid bilayers which is mediated by vesicle adhesion.^70^ Second, both buffers and ions can affect the bending rigidity of the membrane, ^71,72^ but effects depend on the membrane composition.^73^

Strong adhesion is known to trigger pore formation and thus GUV content leakage at least temporarily, which is incompatible while measuring MPs with vectorial transport functions. Temporary leakage also leads to inhomogeneous concentrations of encapsulated dyes used for detection of MP function and complicates quantitative comparison between different vesicles. A limited number of studies has dealt with this topic in detail in the past. Generally, polar and charged molecules seem to have low permeability compared to nonpolar molecules,^43^ but increased permeability for certain low molecular weight compounds has been observed for GUVs compared to LUVs^42^ as well as the co-existence of low and high permeability GUV populations.^43^ Proton permeability is difficult to quantify and the order of magnitude varies greatly in the literature, relating to differences in experimental conditions.^74,75^ The composition of the membrane and the presence of MP further seems to influence the proton permeabilty.^25,76,77^ Using a microfluidic approach, Dimova and colleagues found a slightly lower proton permeability for GUVs compared to LUVs.^25^ Here, we were focused on the leakiness of immobilized GUVs and if immobilization strength can induce leakage. Interestingly, increased leakage of HPTS for GUVs between 7 and 20 µm with strong adhesion was only observed for GUVs prepared by PVA formation but not with Pt wire GUVs. The difference in behavior observed here for PVA and Pt wire GUVs is not obvious, but trace amounts of PVA in or on the membrane can alter membrane property as suggested by Dao et al. 2017.^78^ The difference observed for Pt wire GUVs with and without NaCl is unclear, but could also be due to alteration of membrane properties, such as the bending rigidity, in the presence of salt.^72,73,79^

Proton permeability was tested using GUVs with encapsulated HPTS in the presence of a proton gradient. The results of the HPTS leakage experiments helped us to settle for immobilization conditions that showed immobilization and minimal HPTS leakage. The proton gradient was applied by exterior washing of the GUVs, thus mimicking standard procedures during a biochemical experiment. Only minimal increase in the HPTS ratio was observed during the presence of a proton gradient and most GUVs showed a response to the addition of the protonophore gramicidin, indicating a low level of proton leakage. Our data suggest that immobilization should and can be fine-tuned depending on the experimental set-up. Encouragingly for studies of vectorial proton transport experiments, GUVs between 5 and 20 µm that retain HPTS seem to be also tight against protons.

Different strategies have been described to reconstitute membrane proteins into GUVs. While partial dehydration of deposited membrane protein before lipid rehydration seems to work well for some proteins, a milder procedure might be suitable for large multisubunit complexes such as the members of respiratory chain or eukaryotic proteins. Since reconstitution of many membrane proteins into SUVs are established, transfer from SUVs to GUVs via membrane fusion seems an attractive strategy. We and others have successfully applied charge mediated fusion for the insertion of SUV embedded proteins into the GUV membrane, retaining their functionality, and it is currently the method of choice in our lab and was also applied here.^45,46^ The effect of salt on the fusion of liposomes with oppositely charged lipids has not been conclusively discussed in the literature as some observed no or little influence^45,63^ and others reported impeded fusion in presence of salt.^46^ Using our protocols, we clearly observe rapid and efficient fusion of unlabeled GUVs with fluorescently labeled SUVs both in the presence and absence of salt. However, the fluorescence intensity observed in the GUV membrane is reduced to 50% if fusion is performed in presence of 100 mM NaCl, although the fusion kinetics are similar under both conditions. In fusion experiments with non-immobilized GUVs, PVA GUVs still show a slightly decreased fusion behavior in presence of salt, but the difference was almost gone in PT wire GUVs. We speculate that this might result from different lipid composition of the GUV membrane, i.e. asymmetry of composition in the two leaflets after electroformation^80^ or that trace amounts of PVA might influence the fusion behavior in presence of salt.^78^

Finally, we correlated fusion efficiency with incorporation yield of labelled *bo*_*3*_ oxidase into PVA GUVs without salt. We observed a linear dependence between *bo*_*3*_ oxidase signal and lipid-coupled dye if fusion was performed using labelled proteoliposomes. We concluded that the intensity of lipid-coupled dye correlates well with the intensity of labeled *bo*_*3*_ oxidase. This is an important finding as it allows to estimate protein incorporation without the need of protein labeling. Unfortunately, the method is unable to distinguish between full fusion and hemifusion or simple adhesion of the vesicles to the GUV membrane. While we did not observe a GUV signal increase if neutral SUVs were used, positively charged SUVs might still be able to adhere to vesicles in the presence of salt and undergo lipid mixing without fusion, as suggested by Ishmukhametov et al.^46^ In our previous experiments, however, we were able to show that fusion of SUV and GUV yields functional GUVs and that the fusion behavior was similar to fusion events between oppositely charged SUVs.^45^ A possible solution are content mixing assays in which fusion of both membrane leaflets is a prerequisite to trigger a signal change. However, aside from enzyme-mediated assays, content mixing assays are rather difficult due to unwanted interaction of the often negatively charged cargo with the positively charged lipids. Recently, Lira et al.^63^ described a FRET based assay allowing them to distinguish between hemifusion and full fusion, opening new avenues for verification of complete membrane fusion.

## Concluding remarks

The application of GUVs in a variety of biochemical and biophysical disciplines is very attractive and a broad variety of fascinating results have been described. In this work, we want to contribute to a broader application of GUVs as a system of choice for the investigation of membrane proteins incorporated into the GUV membrane, elevating GUVs from their current main role as sealed lipid container or membrane mimicking system used for docking or membrane deformation studies. We therefore have somewhat systematically analyzed existing methods for GUV formation and immobilization that are accessible without specialized equipment or knowledge and tested relevant properties for membrane protein reconstitution and substrate transport. Although both PVA assisted and electroformation methods used here are based on solvent-free systems, they differ in some of the investigated properties. Electroformation has been the method of choice for many researchers as it is well established and produces clean GUVs. We find that polymer assisted swelling has several advantages, like its ease of use, versatility, scalability, and the production of high GUV numbers, but further research in other materials than PVA and agarose is needed to establish whether polymer assisted swelling can produce GUVs free of impurities.^78,81^ Obviously, powerful alternatives not described here exist, such as microfluidic or oil-emulsion techniques for GUV generation or detergent-mediated reconstitution techniques. There are also efforts to replace natural lipids as used here by more robust synthetic polymers,^25,77,82–84^ and while they surpass GUVs properties in terms of robustness towards mechanical and chemical stresses and life span, they are rather unlikely to match the properties of natural lipids to accommodate for membrane protein thickness and annular lipid layer. GUVs are thus designated to play a critical role also in the future, and robust and widely applicable methods for their generation and use is of importance for all researchers interested.

## Supporting information

Fusion movie

3D printer File - S1

## Acknowledgement

We thank Dr. Thomas Schick for establishing electroformation on ITO coated glass slides and the handling of GUVs after formation. We thank Stephan Berger for establishing PVA assisted swelling in our lab. We thank Sofia Hutter for purification and labeling of the cytochrome *bo*_3_ ubiquinol oxidase. The project was supported from funds of the Swiss National Science Foundation (No. 176154).

## Experimental Procedures

### Materials

1,2-dioleoyl-sn-glycero-3-phosphocholine (DOPC), 1,2-dioleoyl-sn-glycero-3-phospho-(1’-rac-glycerol) (sodium salt) (DOPG) 1,2-dioleoyl-3-trimethylammonium-propane (chloride salt) (DOTAP), 1,2-distearoyl-sn-glycero-3-phosphoethanolamine-N-[biotinyl(polyethylene glycol)-2000] (ammonium salt) (DSPE-PEG(2000) Biotin) and 1,2-dioleoyl-sn-glycero-3-phosphoethanolamine-N-(lissamine rhodamine B sulfonyl) (ammonium salt) (Liss Rhod PE) were obtained from Avanti Polar Lipids (Alabaster, AL, USA), Streptavidin from IBA-Lifesciences (Göttingen, Germany), Invitrogen(tm) 8-Hydroxypyrene-1,3,6-Trisulfonic Acid, Trisodium Salt (HPTS) from Thermo Fisher Scientific (Waltham, Massachusetts, USA), DY-647P1 Maleimide from Dyomics (Jena, Germany) and Polyvinyl alcohol (PVA), fully hydrolyzed, molecular weight approximately 145’000 for synthesis from Merck (Darmstadt, Germany). Other chemicals were obtained from Sigma (St. Louis, Missouri, USA).

### PVA assisted GUV formation

GUV formation with PVA was done as described^49^ with a few modifications. 1 mL 5% PVA (*w/v*) in 200 mM sucrose was incubated for 1 h at 90°C in a thermal shaker lite (VWR international GmbH, Dietikon, Switzerland) at 1’000 rpm, vortexing every 15 – 20 min. A coverglass (25 mm Ø # 1.0, VWR international GmbH) was rinsed with 70% ethanol and placed on an aluminum-foil-covered heat plate set to 50°C. Rubber O-rings with 20 mm diameter and 1.4 mm thickness were placed centrally on top of the coverglass. 200 µL PVA solution was pipetted onto the coverglass area inside the rubber ring and the gel was left to dry for 1 h at 50°C. 20 µL of lipids dissolved in chloroform at 1 mg mL^-1^ composed of 68.8 mol% DOPC, 30 mol% DOPG, 1 mol% Liss Rhod PE and 0.2 mol% DSPE-PEG(2000) Biotin were evenly distributed onto the gels using a 10 µl syringe (MICROLITER™ #701, Hamilton, Bonaduz, Switzerland). Solvent was evaporated for 1.5 – 2 h under vacuum, after which lipids were rehydrated for 1 h using 500 µl formation buffer (5 mM MOPS-BTP pH 7.4, 200 mM sucrose and salt as indicated). The solution was removed from the wells and GUVs were stored for at least 1 h at 4°C before performing further experiments.

### Pt wire GUV electroformation

Electroformation of GUVs using platinum (Pt) wires was performed as described previously^85,57^ with minor modifications. Pt wires were cleaned by hand using soap before incubation in 94% EtOH for 10 min, followed by incubation in chloroform for 10 min, both performed in a sonication bath. Pt wires were dried at RT at 1 atm for 10 min before applying lipid solution to the wires. 10 µL of lipids dissolved in chloroform at 2 mg mL^-1^ with lipid mixture as described were evenly deposited on the wires using a 10µL Hamilton syringe. Lipids were dried on the wires for 1 h in a desiccator under vacuum. For every chamber a coverglass (25mm ø) was coated in a 0.1 g mL^-1^ milk powder solution for 30 min at RT and mild agitation. The coverglass was rinsed with dH_2_O and glued to the formation chamber using Dublisil 22 plus silicone (Dreve Dentamid GmBH, Unna, Germany). Pt wires were connected to an AC electric filed generator (PCGU1000, Velleman Group, Gavere, Belgium) using JST XH2.54 cable with 2 pin female socket (Play-Zone GmbH, Steinhausen, Switzerland) connected to a BNC male to 2 pin terminal block cable (Delock, Berlin, Germany). The formation chamber was filled with 800 µL formation buffer. The following wave sequence protocol was applied: 5 min at 0.44 V_PP_, 5 min at 0.88 V_PP_, 15 min at 1.32 V_PP_, 30 min at 1.76 V_PP_ and overnight at 2.2 V_PP_. Every step was performed at 500 Hz.

### ITO GUV electroformation

Electroformation using indium tin oxide (ITO) coated glass slides was performed as described previously^57,85^ with minor modifications. ITO coated slides were cleaned by hand using soap and incubated in 94% EtOH for 10min in a sonication bath. 10 µL of lipids dissolved in chloroform at 1 mg mL^-1^ with lipid mixture as described were evenly distributed on two separate slides on the conductive surface. The slides were dried for 1 h in a desiccator under vacuum. The electroformation chamber was assembled in a custom 3D printed slide holder (Supplementary File S1) using a 1.00 mm rubber spacer and filled with 600 µL formation buffer. The slides were connected to an AC electric field generator (PCGU1000, Velleman Group, Gavere, Belgium) using crocodile clamps (JYE BNC, Play-Zone GmbH, Steinhausen, Switzerland). The same wave sequence protocol was applied as for Pt wire formation.

### Imaging slide preparation

Imaging of GUVs was performed in an 8 well chambered glass slide (#1.5 high performance cover glass, Cellvis, Mountain View, California, USA). To immobilize GUVs, wells were coated using 200 µL T50-buffer (10 mM Tris-HCL pH 8, 50 mM NaCl) containing 50 µg mL^-1^ biotinylated BSA for 30 min at RT with mild agitation. The solution was replaced by 200 µL T50 buffer containing 10 µg mL^-1^ streptavidin unless stated otherwise and incubated as above. Wells were washed once using 200 µL imaging buffer (5mM MOPS-BTP pH 7.4, 200mM glucose and salt as indicated). A desired amount of GUVs (10 – 100 µL) was loaded into imaging buffer with a final volume of 400 µL and GUVs were left to settle for 1 h prior to imaging. For experiments where immobilization was not required, GUVs were imaged in wells coated using only 200 µL T50-buffer containing 50 µg mL^-1^ BSA for 30 min. Washing and GUV loading was performed as described.

### Image acquisition

Microscopy slides were imaged using an inverted fluorescence microscope (Nikon Ti-2 Eclipse with Crest X-light V2 spinning disk module (disk unit 60 µm), Nikon Europe BV, Amsterdam, Netherlands) with a CFI Plan Fluor 40x oil immersion objective (CFI Plan Fluor 40x/1.30 W.D. 0.24, Nikon Europe BV). Brightfield and fluorescence images were recorded by an Andor Zyla 4.2 Plus USB3 camera in Widefield and Spinning Disk Confocal mode using LED light excitation. HPTS was imaged with excitation at 395 nm and 470 nm and emission at 515 nm using appropriate exciter, emitter, dichroic filter cubes. Liss Rhod PE was imaged with excitation at 550 nm and emission at 595 nm and DY-647P1 with excitation at 640 nm and emission at 698 nm using appropriate exciter, emitter, dichroic filter cubes. Z-stacks were recorded in confocal mode from top to bottom (below the slide surface) with a step size of 1 µm and 72 steps. Time series were recorded as indicated.

### Automatic detection of GUVs

Images in the Liss Rhod PE channel were analyzed in FIJI^86^. For Z-stack acquisitions, two custom macros were used to extract the data. With the first macro a background subtraction is performed on all slices of the Z-stack with a rolling ball radius of 100.0, then an average projection of the stack is made and a bandpass filter is applied to improve separation of nearby vesicles by highlighting the contrast between the background and vesicles using the following settings: filter large set to 40 pixels, filter small set to 5 pixels, suppress stripes set to None, tolerance of direction set to 5 %, with autoscale after filtering and saturation of image when autoscaling enabled. Manual thresholding is performed to create a binary mask separating vesicles and background. The second macro cleans up the mask by performing the binary processes Fill Holes, Erode and Watershed and vesicles are identified using the Analyze Particles function with Size set to 0.75 µm^2^ - Infinity, Circularity set to 0.70 - 1.00 and Display Results, Exclude on Edges and Add to ROI manager enabled. For single images, the use of the first macro was omitted and manual thresholding was directly performed on the image. The second macro was then used as described, with Size set to 19.6 - 314.1 µm^2^ (corresponding to vesicles with diameters between 5 and 20 µm).

### Characterization of GUV formation

For characterization of GUV formations, GUVs were formed as described and 10 µL PVA GUVs or 100 µL electroformed GUVs were loaded to imaging wells coated with BSA as described. 9 Z-stacks were recorded in each well using a 3 × 3 multi spot acquisition from the top left corner of a well to the bottom right corner. GUVs were automatically identified from the Z stack as described. Using the Measure function in FIJI, the Feret diameter of the identified particles is obtained. To estimate the concentration of GUVs with a diameter between 5 – 20 µm, the number of identified particles with the according Feret diameters were counted. Based on the image dimensions (332.8 µm × 332.8 µm), a total area of 0.9968 mm^2^ for all 9 images was obtained which was used to calculate the amount of GUVs per mm^2^. This value was then multiplied by the area of the well which was taken as 80.91 mm^2^, according to the manufacturer, to estimate the amount of GUVs in the entire well chamber. This number was then divided by the volume of GUV solution which was added to the well to obtain the concentration of GUVs (5 – 20 µm) in solution. As a measure for the quality of the formation to produce vesicles with the desired size, the number of particles (5 – 20 µm) was divided by the total number of particles detected as described.

### Calculation of Lipid Yield

To estimate the lipid yield, the total surface area *A*_*GUV*_ of each vesicle was calculated using equation 1 based on the measured Feret diameter *d*_*GUV*_ and the bilayer thickness *d*_*bilayer*_ which was assumed to be 5 nm.

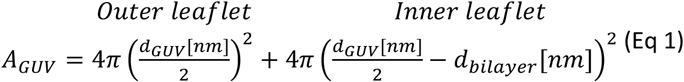

The lipid yield was calculated using equation 2.

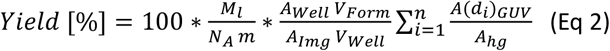

The total number of lipids in GUV *i* with a diameter of *d*_*i*_ was calculated by dividing the surface *A(d*_*i*_)_*GUV*_ calculated according to Eq 1 by the surface area of the lipid headgroup *A*_*hg*_ which was assumed to be 0.71 nm^2^.^87,88^ The total number of lipids from all GUVs was calculated by summation of the lipid number of each GUV *i* with *n* as the total number of particles detected in one formation condition. To estimate the total number of lipids in the formation solution after removal from the formation chamber, the sum was divided by the total imaged area *A*_*Img*_ of the 9 images (0.9968 mm^2^) times the volume *V*_*Well*_ of GUV solution added to the well (for example 10 µL for PVA GUVs) and multiplied with the total well area *A*_*Well*_ (80.91 mm^2^) times the volume *V*_*Form*_ of solution used for the formation of GUVs (for example 500 µL for PVA GUVs). The yield was then obtained by dividing the total number of lipids in the formation solution by the Avogadro constant *N*_*A*_, multiplying with the average molecular weight *M*_*l*_ of the lipids which was assumed to be 790 g mol^-1^, based on a lipid composition of roughly 70 mol% DOPC and 30 mol% DOPG, and finally divided by the lipid weight *m* used to coat the surface for GUV formation (20 µg). This was multiplied by 100 to obtain values in percentage.

### Immobilization assay

Flow immobilization experiments were performed in channel slides (µ-Slide III 3D Perfusion uncoated, ibidi GmbH, Gräfelfing, Germany) connected to a syringe pump (Ossila, Sheffield, UK). All experiments were performed at a flow rate of 5 µL s^-1^. The slide was coated as described using 30 µL 100 µg mL^-1^ biotinylated BSA followed by 30 µL 0, 10 or 100 µg mL^-1^ streptavidin in T50 buffer. 1 µL 0 mM NaCl and 5 µL 100 mM NaCl PVA GUVs were loaded into imaging buffer to a final volume of 30 µL and GUVs were left to settle for 1 h. Prior to imaging the slide was sealed and the channels were filled with imaging buffer according to the manufacturer protocol. Image acquisition was performed using the confocal Liss Rhod PE settings and a time series was recorded over 2 min with a 0.5 s interval and 100 ms exposure at 50% laser intensity. Flow was started after 15 s using the syringe pump. GUVs were automatically detected as described on the last image before visible flow. To detect non-immobilized vesicles, all images recorded during flow were transformed into an average Z-projection, resulting in smearing and a decreased intensity for non-immobilized GUVs. This allowed automatic detection of immobilized GUVs by thresholding as described. The percentage of immobilized GUVs was calculated as the ratio between the detected number of GUVs on the average projection and before flow.

### Calculation of the streptavidin density

To achieve equal slide coating, the streptavidin amount per surface was calculated based on the microscopy well dimensions indicated by the slide manufacturers. For simplicity reasons, we will refer to this value as the streptavidin density. For 8 well chambered slides, well dimensions are 8.7 × 9.3 mm leading to a bottom surface of 80.91 mm^2^. Based on the coating volume of 200 µL, the height of the coated wall was estimated to be 2.47 mm, giving a total wall surface of 88.92 mm^2^ leading to approximately 169.83 mm^2^ of surface that is coated. This gives a volume to surface ratio of 1.18 µL mm^-2^. For ibidi slides, the well diameter is 5.5 mm with a well height including the channels of 1.7 mm. This gives a bottom surface of 23.75 mm^2^ and a wall surface of 29.37 mm^2^. The well is connected by two channels which have a width of 1 mm and a height of 0.5 mm, which takes approximately 1 mm^2^ away from the wall surface. This gives a total surface of 52.12 mm^2^ and a volume to surface ratio of 0.58 µL mm^-2^ with 30 µL coating volume which is approximately half of the ratio for the 8 well chambered slide. Thus, to get similar immobilization conditions for both slides, the concentration of streptavidin for the ibidi slides should be twice as high as for the 8 well chambered slide.

### HPTS leakage

For PVA GUVs, HPTS leakage during immobilization was measured at streptavidin densities of 0, 6, 30 and 60 ng mm^-2^ streptavidin as calculated above. HPTS was either added to imaging buffer before GUV addition or after immobilization. Final HPTS concentrations in the well chamber were 43 - 50 µM. 8 well chambered slides were prepared as described and 390 µL or 350 µL appropriate imaging buffer containing HPTS and 10 µL or 50 µL 0 mM NaCl or 100 mM NaCl GUVs were loaded, respectively. For leakage after immobilization, 375 µL appropriate imaging buffer as well as 25 µL GUV solution was loaded, whereby 5 µL 0 mM NaCl GUVs were diluted with 20 µL formation buffer and 100 mM NaCl were used undiluted. GUVs were left to settle, after which 100 µL appropriate imaging buffer containing 230 µM HPTS was added and incubated for 1h before imaging. For Pt wire GUVs, HPTS leakage during immobilization was measured at coating densities of 6 and 60 ng mm^-2^ streptavidin. 8 well chambered slides were prepared as described and 380 µL or 350 µL appropriate imaging buffer containing HPTS and 20 µL or 50 µL 0 mM NaCl or 100 mM NaCl GUVs were loaded, respectively. GUVs were imaged by recording Z-stacks using the Liss Rhod PE channel and the 470 nm excitation channel for HPTS.

### HPTS leakage analysis

GUVs were automatically detected from Z-stacks as described. The number of GUVs considered for analysis was limited to 148 for each sample. 20 background regions of interest (bgROIs) were drawn in by hand in areas that did not contain GUVs. For leakage analysis, a background subtraction with a rolling ball radius of 200 pixels and light background enabled was first performed on each slice of the Z-stack in the HPTS channel. An intensity profile of the GUVs and bgROIs in the Z dimension was recorded using the Time Series Analyzer V3 plugin in FIJI with the average intensity setting. The profiles were normalized to the average of the first 20 steps of the Z-stack. The profiles of the bgROIs were averaged to create a mean background profile. This was then subtracted from each GUV profile which were then inverted by multiplying with -1 to obtain positive values for GUVs that did not leak. The maximum intensity of the profile was then measured, indicating the largest absence of HPTS signal for each GUV. Only GUVs in the desired size range (5 – 20 µm diameter) were used for analysis. Because small GUVs tended to move at low streptavidin concentrations and were only imaged by very few slices, vesicles with diameters below 7 µm were further discarded as well. We further limited the number of vesicles from each experiment to 85 due to limitations in GraphPad Prism 8.0 which resulted in 20 – 85 vesicles that were analyzed. The median of the maximum intensity distribution of the GUVs was calculated in Prism and used to define a threshold for leakage based on the average median at low streptavidin concentrations. Leaky GUVs were defined by having a maximum intensity below approximately 25% of the average median. The percentage of leaky GUVs was calculated by dividing the number of GUVs below the threshold by the total number of GUVs.

### Proton leakage

For proton leakage, formation buffers contained 10 mM MOPS-KOH pH 7.4, 200 mM sucrose and imaging buffers were 10 mM MOPS-KOH pH 7.4, 200 mM glucose and 10 mM MOPS-KOH pH 8.0, 200 mM glucose. Buffers for GUVs with salt contained 100 mM NaCl as well. 8 well chambered slides for immobilization were prepared as descried. For PVA GUVs, 400 µL imaging buffer and 20 µL 0 mM NaCl or 50 µL 100 mM NaCl GUVs were loaded, respectively. For Pt wire GUVs, 400 µL imaging buffer and 50 µL GUVs were loaded, both for 0 mM NaCl GUVs and 100 mM NaCl GUVs. Before imaging, a pH exchange in the exterior solution was performed by performing two 1 mL wash steps with imaging buffer at pH 7.4 followed by two 1 ml wash steps with buffer at pH 8.0. Washing was performed using two 1 mL pipettes with simultaneous addition and removal of solution in a well chamber. A single image was acquired in confocal mode using the Liss Rhod PE channel and GUVs were then recorded in confocal mode using both HPTS channels for 60 min at 5 min intervals. Approximately 1 min after the 60 min acquisition (61 min), another time series was recorded for 5 min and 5 s intervals with addition of 5 µL 1 mM gramicidin (final concentration 12.5 µM) after 1 min (62 min). The total acquisition time was 66 min.

### Proton leakage analysis

GUVs were identified using the single Liss Rhod PE image as described. The HPTS intensity for both channels were extracted using the Time Series Analyzer V3 plugin in FIJI with the average intensity setting. The HPTS ratio was calculated by dividing the intensity at 470 nm excitation by the intensity at 395 nm excitation. To assess the amount of leakage after 20 min, the ratio at 0 min was subtracted from the ratio at 20 min. As we expect an increase in pH on the inside as GUVs leak, only vesicles with a ratio difference > 0 were considered for further analysis. The ratio at 66 min was then subtracted from the ratio at 0 min, indicating the total possible increase for a GUV. The ratio difference 20 min – 0 min was then divided by the total possible increase and multiplied with 100 to obtain a percentage of leakage.

### Preparation of SUVs

Liposomes were formed with a lipid composition of 69 mol% DOPC, 30 mol% DOTAP and 1 mol% Liss Rhod PE or 99 mol% DOPC and 1 mol% Liss Rhod PE. Lipids dissolved in chloroform were mixed in a 25-mL round bottom flask and chloroform was evaporated under a constant stream of N_2_ while rotating the flask. The film was further dried overnight in a desiccator under vacuum. Lipids were resuspended in formation buffer to a concentration of 5 mg mL^-1^ and unilamellar vesicles were obtained by performing seven freeze-thaw cycles and stored at -80°C before further use. For fusion experiments, liposomes were thawed and diluted to 50 µg mL^-1^ using imaging buffer and size was adjusted by sonication on ice using a tip sonicator (Vibra Cell 75186, Thermo Fisher Scientific, Waltham USA) for 2 min with 30 s ON and 30 s OFF pulses and 40% amplitude.

### Cytochrome *bo*_*3*_ ubiquinol oxidase purification and labeling

Cytochrome *bo*_*3*_ ubiquinol oxidase mutant IIIA21C, encoded by a cysless pETcyoII plasmid, was expressed and purified as described.^89–91^ The protein was cysteine-labeled with DY-647P1 maleimide as previously published^92^ with minor modifications. The excess dye was removed by performing a CentriPure P50 (emp BIOTECH, Berlin, Deutschland) size exclusion chromatography followed by a Superdex 200 Increase 10/300 GL column (ÄKTA Pure system, GE Healthcare, Boston, Massachusetts, USA) purification at 4 °C. The labeled protein was concentrated to 20 µM.

### Reconstitution of cytochrome *bo*_*3*_ ubiquinol oxidase

DOTAP liposomes were formed and sonicated at 5 mg mL^-1^ as described. 240 µL liposomes were destabilized using a final concentration of 0.4% (*w/v*) sodium cholate, mixed with 12 µL purified *bo*_*3*_ oxidase (20 µM) and incubated on ice for 30 min to obtain proteoliposomes. Empty liposomes were prepared by addition of 12 µL buffer instead of protein. Liposomes were collected by gel filtration using a CentriPure P10 column (*emp* BIOTECH GmbH, Berlin, Germany) pre-equilibrated with formation buffer. The column was eluted using 1.2 mL formation buffer to obtain 1 mg mL^-1^ liposomes. Prior to measuring, the liposomes were diluted to 0.2 mg mL^-1^ using formation buffer and proteoliposomes and empty liposomes were mixed in a 3/0, 2/1, 1/2 and 0/3 ratio.

### Charge mediated fusion

Charge mediated fusion of negatively charged GUVs and positively charged SUVs was performed in 8 well chambered slides and in 1.5 mL Eppendorf tubes. GUVs were prepared as described and 10 – 50 µL GUV solution was diluted in formation buffer to a final volume of 50 µL to achieve similar vesicle concentrations. For fusion in an 8 well chambered slide, diluted GUVs were loaded into imaging buffer containing salt as indicated to a final volume of 400 µL and immobilized for 1 h. A time series was recorded for 5 min at 5 s intervals using confocal Liss Rhod PE settings for all experiments and additional DY-647P1 settings for experiments with proteoliposomes. Fusion was initiated by addition of 100 µL SUVs after 30 s. Final concentration of SUVs was 10 µg mL^-1^ for experiments with PVA and Pt wire GUVs and empty vesicles and 40 µg mL^-1^ for experiments with PVA GUVs and mixtures of empty and proteoliposomes. For fusion in Eppendorf tubes, diluted GUVs were mixed with 50 µl empty SUVs (20 µg mL^-1^) and incubated for 15min at RT. 100 µL of fused GUVs were transferred into 400 µL imaging buffer and GUVs were immobilized for 1 h. Z-stacks were recorded as described using confocal Liss Rhod PE settings.

### Analysis of charge mediated fusion

For fusion of PVA and Pt wire GUVs with empty SUVs in 8 well chambered slides, 10 – 50 GUVs were detected as described using a single Liss Rhod PE image recorded after 300 s. A bgROI was manually drawn next to every vesicle. The Liss Rhod PE intensity was extracted for every GUV and corresponding bgROI using the Time Series Analyzer V3 plugin in FIJI with the average intensity setting. For each GUV, the background was removed by subtracting the intensity of the corresponding bgROI. The mean intensity increase of GUVs per experiment was calculated and average and standard deviation of mean intensity values from four replicates were plotted. To show end point signal distributions, the final Liss Rhod PE intensity of individual GUVs of the four replicates were combined and plotted. For experiments using proteoliposomes, 30 – 50 GUVs were detected as described after 180 s. On the same image, mean grey intensity for Liss Rhod PE and DY-647P1 of GUVs and corresponding bgROIs were extracted using the Measure function of FIJI. Background was subtracted as described. To show signal correlation, the Liss Rhod PE signal of every GUV was plotted on the X-axis against the DY-647P1 signal of every GUV on the Y-axis. For fusion experiments in Eppendorf tubes, GUVs were automatically identified from Z-stacks as described. An average Z-projection was performed and the background was subtracted using a rolling ball radius of 200 pixels. Using the Measure function in FIJI, the mean grey intensity of the Liss Rhod PE channel was extracted for every GUV and plotted as a signal distribution.

## Notes

### Competing Interest Statement

The authors have declared no competing interest.

