## Supplementary Information for "Experimental platform for the functional investigation of membrane proteins in giant unilamellar vesicles"

* Contributed equally to the manuscript

### Supplementary Results and Discussion

**Lipid yield and average diameter**

To estimate the molar yield (based on deposited lipid) and the average diameter of giant unilamellar vesicles (GUVs), we used the size distribution data and the GUV concentration obtained from each experiment as described in the Materials and Methods. The mean and standard deviation of the molar yield of free-standing GUVs and average diameter are presented in Table S1. Not surprisingly, the highest yield (~30% of deposited lipid was incorporated into GUVs) was obtained for PVA formation in the absence of salt which is similar to what has recently been reported for polymer assisted swelling using cellulose.[65] In the presence of 100 mM monovalent salt, the yield decreased to 17 - 19% for PVA formation. All electroformations produced similar yields with around 4% under the experimental conditions used here. Unlike PVA formation, the yield was not affected by the presence of monovalent salt although a slight decrease in GUV number was observed (from ~6 x$10^5$ GUVs $mL^{-1}$ in absence to ~3-4 x$10^5$ GUVs $mL^{-1}$ in presence of salt in the requested size). This is explained by the larger average diameter of salt GUVs that require more lipids per GUV. Thus, the decreased concentration is balanced by the increase in frequency of large Pt wire GUVs in the presence of salt, resulting in similar yields for those formations. The increased average diameter reflects the increase in frequency of large GUVs (> 10 µm) for salt GUVs (Figure S3 and S4).

**HPTS leakage**

Soluble fluorescent dyes are frequently used to detect substrates translocated across the lipid bilayer by membrane proteins (MPs). These dyes are usually encapsulated in GUVs and need to remain inside the vesicle for the duration of the experiment. Therefore, leakage of the dye, for example due to strong vesicle adhesion, would be detrimental for measuring MP activity. Thus, we investigated whether GUVs become permeable to the dye HPTS due to immobilization. After vesicles were left to immobilize, they were imaged by recording a Z-stack using both the Liss Rhod PE and HPTS channels. The signal from the labeled lipid Liss Rhod PE was used to identify the GUV membrane while leakage was assessed using the HPTS signal. For each vesicle, an average signal for each Z-slice was obtained which was normalized and inverted to obtain values close to 0 for vesicles with HPTS leakage and positive values for vesicles without leakage (see Material and Methods for the exact process). To distinguish between leaky and non-leaky GUVs (Figure S6A), the maximum intensity value from the whole Z-stack (max Z intensity) was used for each vesicle and a threshold using the average median value (0.006) from GUVs at weak adhesion was defined. GUVs were thus denominated as leaky if the max Z intensity was below 25% of the median value (<0.0015). As small vesicles (< 7 µm) were recorded by only few slices and tended to be less well immobilized, they were discarded from the analysis, as were large GUVs > 20 µm which are outside of our desired size range. Figure S6B shows the distribution of max Z intensity values for PVA and Pt wire GUVs with and without 100 mM sodium chloride. Although slight shifts in the distribution are observed for all GUVs at high streptavidin densities, only PVA GUVs show a clear increase in the number of vesicles with values below the threshold at high streptavidin density, indicating increased leakage for PVA GUVs with strong immobilization. For Pt wire GUVs, only the number of fully leaky GUVs was slightly higher without salt compared to 100 mM NaCl and occasional HPTS leakage at strong adhesion was only observed in Pt wire GUVs larger than 20 µm. Percentage of leaky GUVs for each of the three replicates (Figure S6C) were calculated. An increase in leakage at high streptavidin density is again observed for PVA GUVs, although the error is quite large at high streptavidin density. Importantly, even at high streptavidin concentration, the percentage of leaky GUVs never exceeded 35%. Interestingly, GUVs formed with the Pt wire method were less affected by the increased streptavidin concentration compared to PVA GUVs. Generally, the number of leaky GUVs at weak adhesion was low.

Because we observed large differences in the number of leaky GUVs for PVA assisted formation at 6 and 60 ng mm$^{-2}$ streptavidin, we repeated the experiment with 0, 6, 30 and 60 ng mm$^{-2}$ streptavidin (Figure S7A and B). Increased leakage of HPTS to the inside of GUVs was already observed at 30 ng mm$^{-2}$ streptavidin. However, at 6 ng mm$^{-2}$ where low leakage is still observed, PVA GUVs with no salt were poorly immobilized (20% as opposed to 100% with sodium chloride), thus a streptavidin density between 6 and 30 ng mm$^{-2}$ should be used. In the previous experiment, we were not able to discriminate if leakage happens only during immobilization and tightness is restored or if the leakiness is permanent. This is critical, as substrates or other chemicals are added after the immobilization process. We focused on PVA GUVs due to the observed differences at various streptavidin densities and added solution containing HPTS to vesicles that were already immobilized on the slide (Figure S7C and D). Strikingly, while leakage remained low for PVA GUVs without salt (< 10%), it was highly elevated for PVA GUVs with 100 mM NaCl at strong adhesion with more than half of the vesicles showing HPTS influx. Sample images from all these different conditions are depicted in Figure S8.

The two different experiments (leakage during and after immobilization) show a different behavior for PVA GUVs. While no difference was observed for GUVs with and without salt for leakage during immobilization, a large difference was seen when HPTS was added to GUVs that were already immobilized, indicating that PVA GUVs in the absence of salt mostly form pores during the immobilization process while PVA GUVs in the presence of salt remain leaky after immobilization. It is important to note that in the experiment with HTPS addition after immobilization, flow stress is exerted to the vesicles that could lead to increased HPTS permeability, especially if vesicles are unstable.

**Proton leakage**

One of the main interests in our group are MPs form the respiratory chain which translocate protons. Thus, GUVs should be able to maintain a proton gradient. To detect proton leakage, GUVs containing HPTS were prepared at pH 7.4, immobilized and washed with buffer at pH 8.0, resulting in an inside acidic gradient of 0.6. The HPTS signal was monitored for 1 h before the pH gradient was equilibrated by the addition of the protonophore gramicidin, leading to an alkalinization of the GUV lumen and an increase in HPTS ratio. The signal increase of all GUVs that were selected for analysis showed minimal increase of HPTS ratio during the 1 h incubation, which is an indication of GUV tightness towards protons (Figure S10). Some GUVs displayed very small or higher percentages. The latter might indicate high leakage, but we also observed some GUVs which showed only small increase after gramicidin addition, which would in turn result in a large percentage. We are not sure why these GUVs did not react to gramicidin, it is possible that they were multilamellar or that the concentration of gramicidin was not high enough to achieve efficient gramicidin incorporation in all GUVs. Further, some GUVs seem to even decrease in signal after gramicidin addition, potentially due to HPTS leakage, which might even result in negative percentages and most likely account for the outliers at very small values.

**Charge-mediated fusion in an Eppendorf tube**

The last step towards measurements of MPs in GUVs is the functional reconstitution of the protein. Here, we used charge-mediated fusion of positively charged small unilamellar vesicles (SUVs) to negatively charged GUVs as a method to incorporate MPs into GUVs. We started by performing experiments with empty liposomes, initiating fusion directly on the microscopy slide by addition of SUVs to immobilized GUVs. We observed a large distribution of signals, potentially due to effects of immobilization or mixing and pipetting artifacts. To avoid these artifacts, we repeated the experiments by incubation of free-floating GUVs (approximately 0.3 – 1.6 x$10^6$ 5 – 20 µm GUVs per mL) with SUVs for 15 min in an Eppendorf tube after which they were transferred into a microscopy well and left to immobilize. Z-stacks were recorded for all conditions and an average Z-projection was created for every stack. Average signal intensities for every GUV were then extracted and depicted as a signal distribution (Figure S13). Contrary to our expectations, fusion in an Eppendorf tube did not lead to a smaller signal distribution, highlighting that the fusion process is not homogenous. For PVA GUVs, we observe a decrease in signal in the presence of 100 mM NaCl both for fusion in the microscopy well and in the Eppendorf tube. Interestingly, this was not the case for Pt wire GUVs where this trend was not observed for fusion in the Eppendorf tube.

#### Supplementary Tables and Figures

**Table S1: Lipid Yield and mean average diameter of GUVs prepared by PVA formation or electroformation.** The lipid yield was calculated as described in Material and Methods and the mean and standard deviation of three experiments is shown. The average diameter was calculated from the diameter distributions recorded for each formation and the mean and standard deviation of the average diameters for the three experiments are shown. Values could not be determined for formations that produced insufficient GUV numbers and are labeled with ND.

| | PVA | | Pt wire | | ITO | |
|---|---|---|---|---|---|---|
| | Yield [%] | Diameter [µm] | Yield [%] | Diameter [µm] | Yield [%] | Diameter [µm] |
| 0 mM salt | 31.96 ± 12.67 | 3.82 ± 0.16 | 3.69 ± 1.14 | 4.21 ± 0.39 | 4.29 ± 2.66 | 6.40 ± 1.36 |
| 100 mM NaCl | 16.65 ± 1.86 | 3.81 ± 0.33 | 4.18 ± 0.89 | 7.49 ± 0.84 | ND ND | ND ND |
| 100 mM KCl | 18.63 ± 9.10 | 3.94 ± 0.46 | 4.25 ± 0.46 | 6.48 ± 1.09 | ND ND | ND ND |

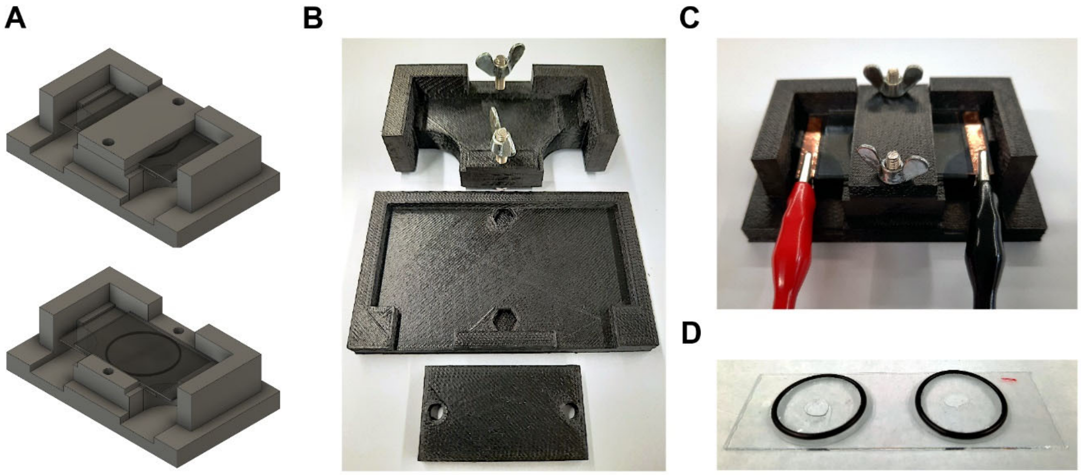


**Figure S1: Assembly of the ITO electroformation and PVA formation chambers.** A) 3D render of the custom ITO-coated slide holder (Supplementary File S1) with (top) and without (bottom) lid to hold the slides in place. Slides are shown with rubber O-ring as spacer. B) 3D printed parts of the slide holder. Holes are designed to fit M4 hex bolts to screw on the lid using wing nuts (not 3D printed). The holder is composed of the main holder (top), a stand (middle) and a lid (bottom). C) Assembled ITO formation chamber. ITO-coated slides are modified with adhesive copper tape and separated by a 1 mm rubber spacer. Crocodile claps are attached to the copper tape. D) PVA formation chamber. Two coverglasses (25 mm Ø) with PVA gel inside a rubber O-ring are glued to a microscopy slide for better transportation using grease for laboratories. The O-ring forms a barrier, allowing the addition of formation buffer to the gels. To protect the formation from dust, slides are covered by a lid (not shown).

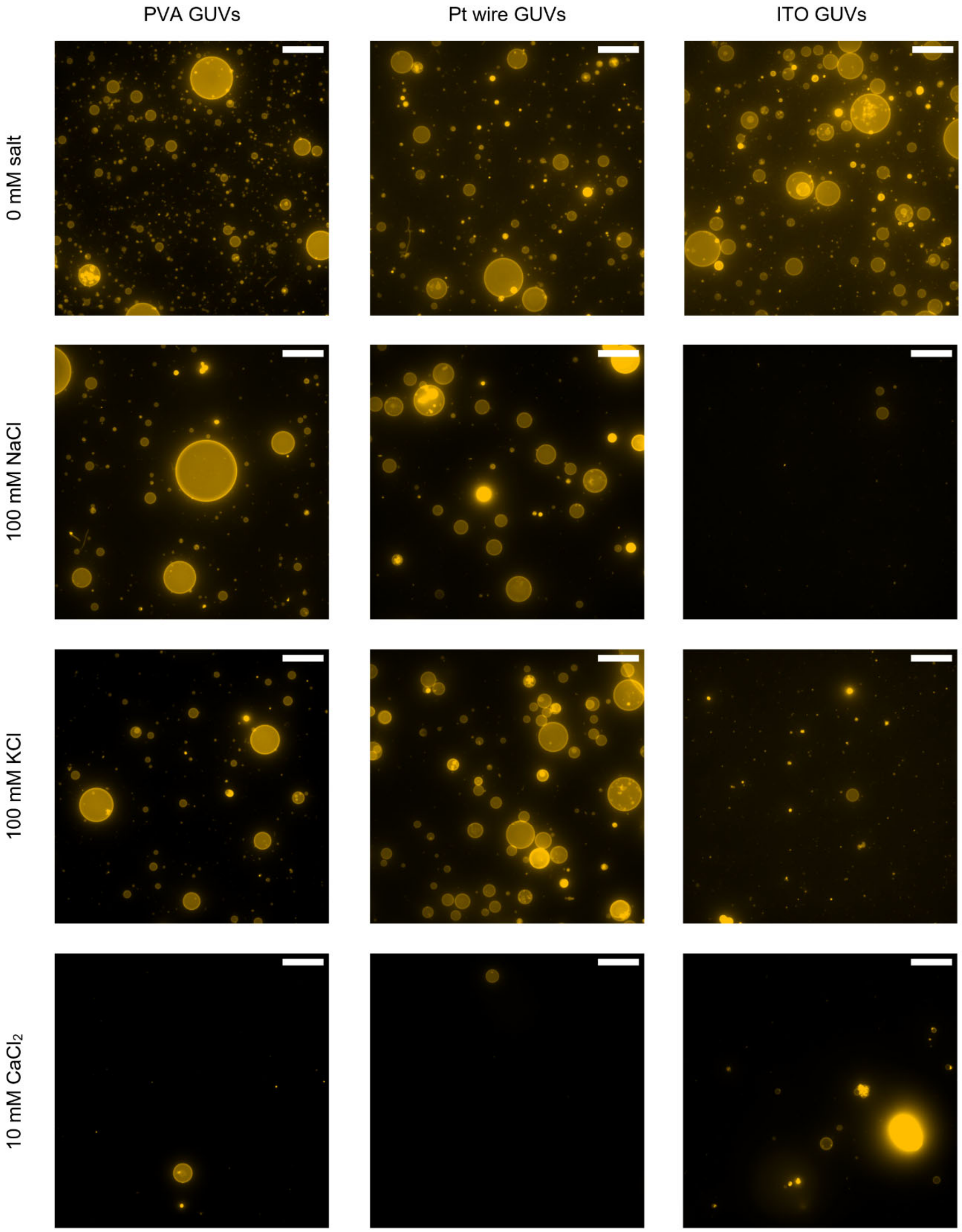


**Figure S2: Representative GUV formations in the presence or absence of salt.** Average Z-projections of confocal Z-stacks recorded in the Liss Rhod PE channel are shown. PVA GUV formations are shown in the first column, Pt wire formations in the second column and ITO formations in the third column. The first row depicts formations in the absence of salt, the second row with addition of 100 mM NaCl, the third row with addition of 100 mM KCl and the last row with addition of 10 mM $CaCl_2$ to the formation buffer. The scale bar is 50 µm.

Here we show the calculated histograms from the size distributions of each formation condition. A bin size of 1 µm was chosen to illustrate the size distribution in the relevant size range of 0 – 20 µm. A larger bin size of 5 µm was chosen to represent the entire size range of the formations (up to 100 µm) and the frequency is displayed in the logarithmic scale, due to the very small frequency of large GUVs.

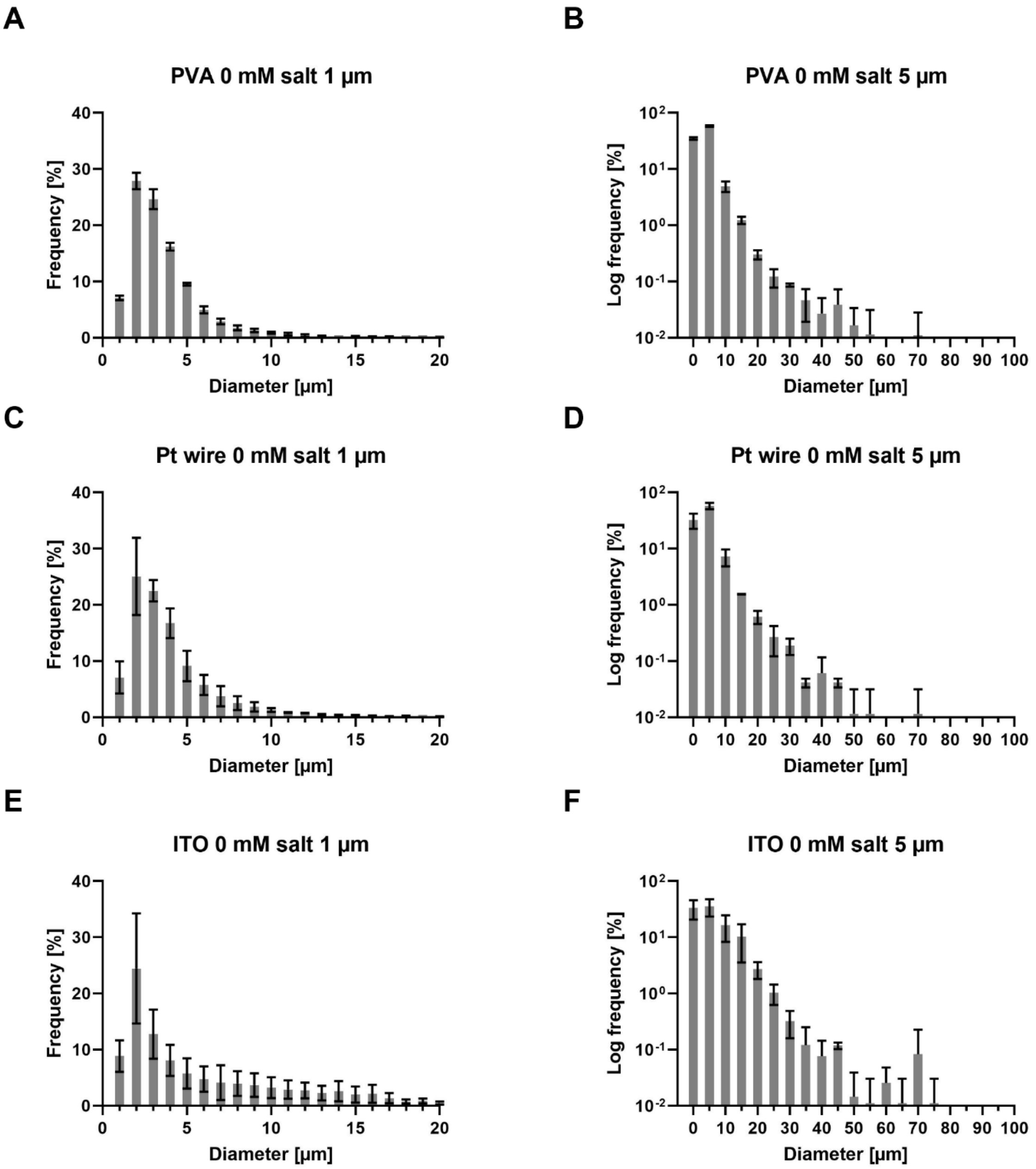


**Figure S3: Histograms of GUV formations in the absence of salt. The bar height represents the average frequency and error bars the standard deviation from three experiments.** A, C, E show histograms with a bin size of 1 µm cut off at 20 µm. B, D, F show histograms with a bin size of 5 µm cut off at 100 µm. The Y-axis is logarithmic as large GUVs tend to have small frequencies. A-B) Histograms from PVA formation. C-D) Histograms from Pt wire formation. E-F) Histograms from ITO formation.

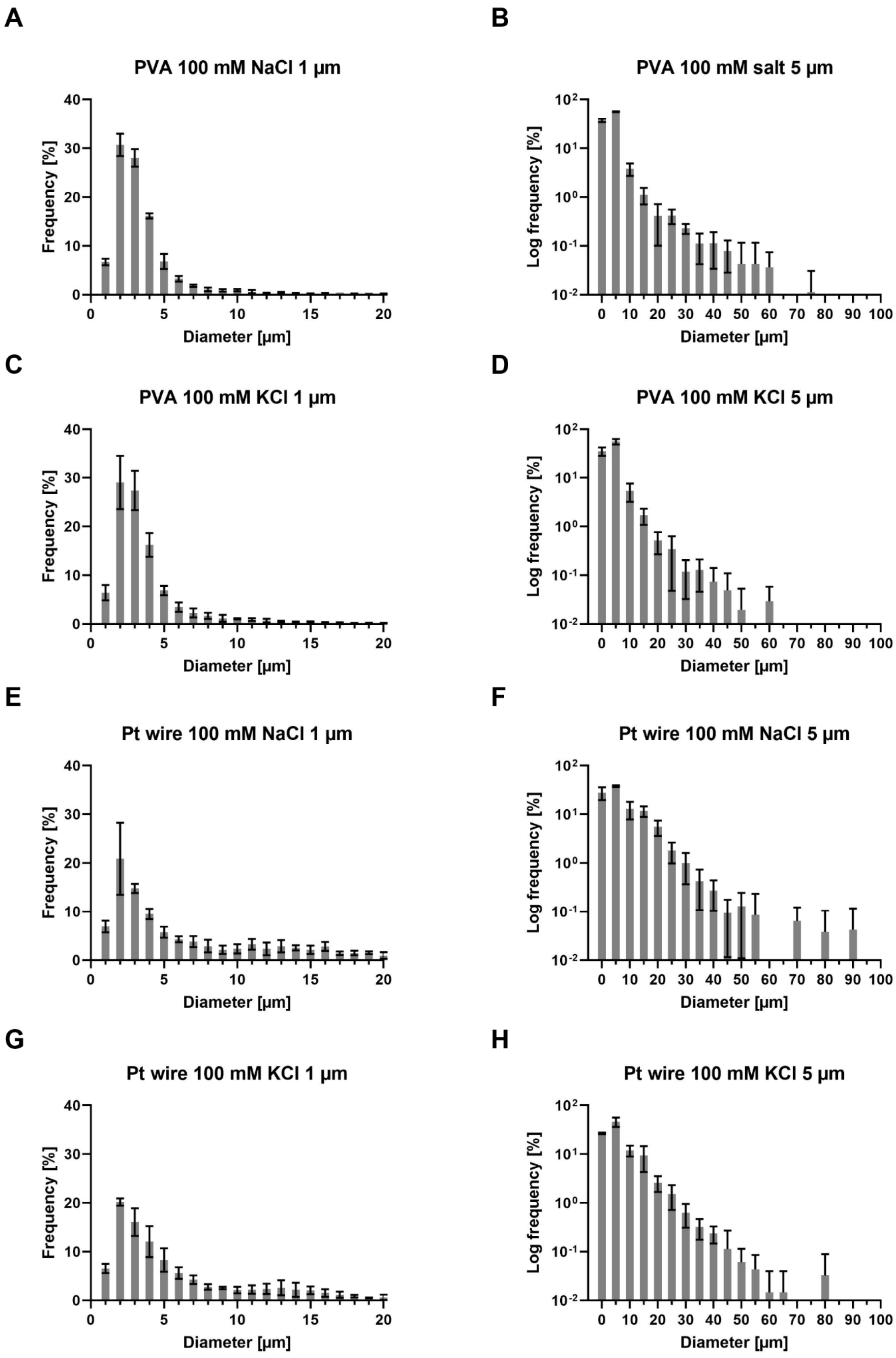
A
PVA 100 mM NaCl 1 µm
Frequency [%]
Diameter [µm]
B
PVA 100 mM salt 5 µm
Log frequency [%]
Diameter [µm]
C
PVA 100 mM KCl 1 µm
Frequency [%]
Diameter [µm]
D
PVA 100 mM KCl 5 µm
Log frequency [%]
Diameter [µm]
E
Pt wire 100 mM NaCl 1 µm
Frequency [%]
Diameter [µm]
F
Pt wire 100 mM NaCl 5 µm
Log frequency [%]
Diameter [µm]
G
Pt wire 100 mM KCl 1 µm
Frequency [%]
Diameter [µm]
H
Pt wire 100 mM KCl 5 µm
Log frequency [%]
Diameter [µm]

**Figure S4 (previous page): Histograms of PVA and Pt wire GUV formation in the presence of salt.** The bar height represents the average frequency and error bars the standard deviation from three experiments. A, C, E, G show histograms with a bin width of 1 µm cut off at 20 µm. B, D, F, H show histograms with a bin width of 5 µm cut off at 100 µm. The Y-axis is logarithmic as large GUVs tend to have small frequencies. A-B) Histograms from PVA formation with 100 mM NaCl. C-D) Histograms from PVA formation with 100 mM KCl. E-F) Histograms from Pt wire formation with 100 mM NaCl. G-H) Histograms from Pt wire formations with 100 mM KCl.

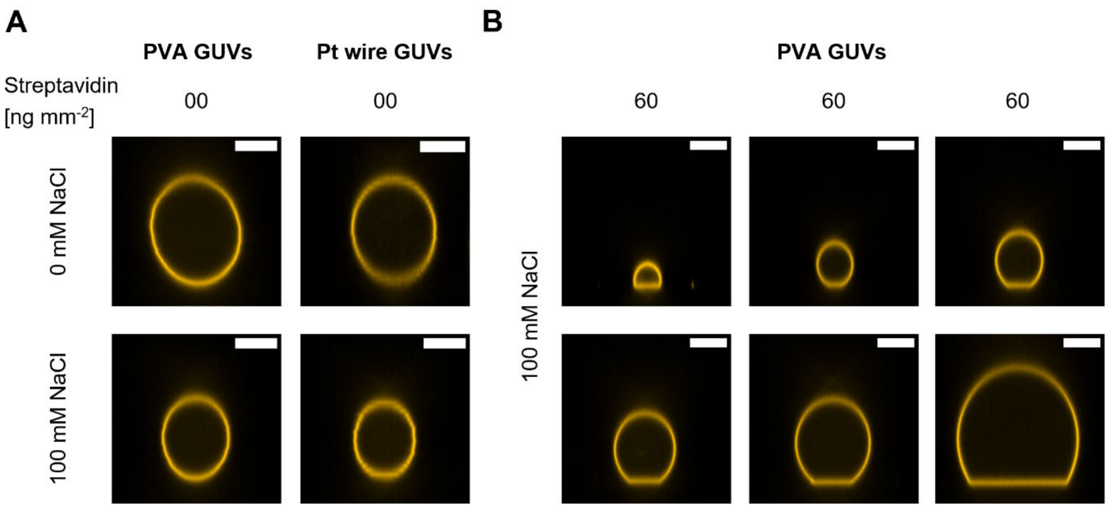


**Figure S5: Spherical adhesion caps in the absence of streptavidin and heterogeneity of caps in a single well.** Confocal Z-stacks were recorded and side-views of representative GUVs are shown. The scale bar is 10 µm. A) PVA and Pt wire GUVs in the presence and absence of 100 mM NaCl were immobilized at a streptavidin density of 0 ng mm$^{-2}$ No Spherical caps are observed. B) Different PVA GUVs in the same well chamber with 100 mM NaCl immobilized at a streptavidin density of 60 ng mm$^{-2}$ are shown.

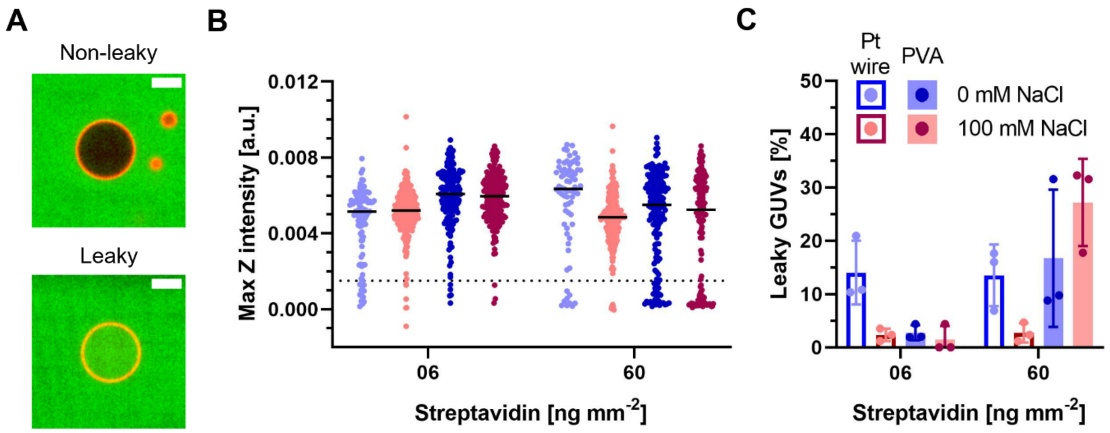


**Figure S6: Immobilization-induced HPTS leakage of PVA and Pt wire GUVs (7 – 20 µm) in the presence (red) or absence (blue) of 100 mM NaCl at low and high streptavidin densities.** A) Example GUV without (top) and with (bottom) HPTS leakage. Overlay of HPTS channel (green) and Liss Rhod PE channel (red). The scale bar is 10 µm. B) Combined distribution of the maximum Z intensity values from three experiments. Each dot represents a single GUV and vesicles from three replicates are combined. The median is indicated by a black line and the threshold is depicted by a dotted line. Maximum Z intensities were extracted from Z stacks recorded in the HPTS channel according to Material and Methods. Values close to zero indicate HPTS leakage into GUVs and positive values indicate the absence of HPTS in the GUV lumen. The dotted line indicates the threshold (0.0015) used to calculate the percentage of leaky GUVs and is approximately 25% of the average median (black lines) of GUVs immobilized at a low streptavidin density. GUVs below the threshold were classified as leaky. C) Percentage of leaky GUVs from B. The height of the bar indicates the average percentage of leaky GUVs with individual values from the three experiments shown as dots. Error bars indicate the standard deviation.

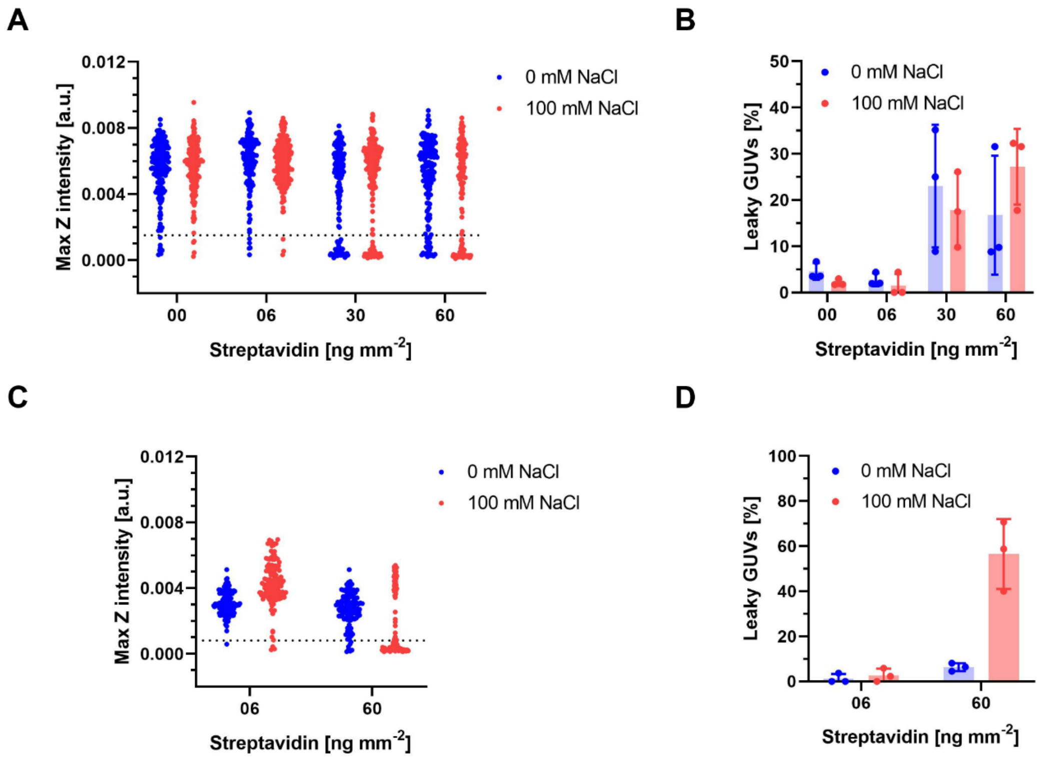


**Figure S7: HPTS leakage of PVA GUVs (7 – 20 µm) in the presence (red) or absence (blue) of 100 mM NaCl at different streptavidin densities.** A and B show immobilization-induced HPTS leakage at 00, 06, 30 and 60 ng mm$^{-2}$ streptavidin. C and D show leakage of HPTS with addition of dye after immobilization at low and high streptavidin density. A and C show the combined distribution of the maximum Z intensity values from three experiments. Maximum Z intensities were extracted from Z stacks recorded in the HPTS channel according to Material and Methods. Values close to zero indicate HPTS leakage into GUVs and positive values indicate the absence of HPTS in the GUV lumen. The dotted line indicates the threshold used to calculate the percentage of leaky GUVs and is approximately 25% of the average median of GUVs immobilized at a low streptavidin densities. GUVs below the threshold were classified as leaky. A) Maximum Z intensity distribution. The threshold (0.0015) was set based on the average median at streptavidin densities of 00 and 06 ng mm$^{-2}$. B) Percentage of leaky GUVs from A. The height of the bar indicates the average percentage of leaky GUVs with individual values from the three experiments shown as dots. Error bars indicate the standard deviation. C) Maximum Z intensity distribution. The threshold (0.0008) was set based on the average median at a streptavidin density of 06 ng mm$^{-2}$. D) Percentage of leaky GUVs from C. The height of the bar indicates the average percentage of leaky GUVs with individual values from the three experiments shown as dots. Error bars indicate the standard deviation.

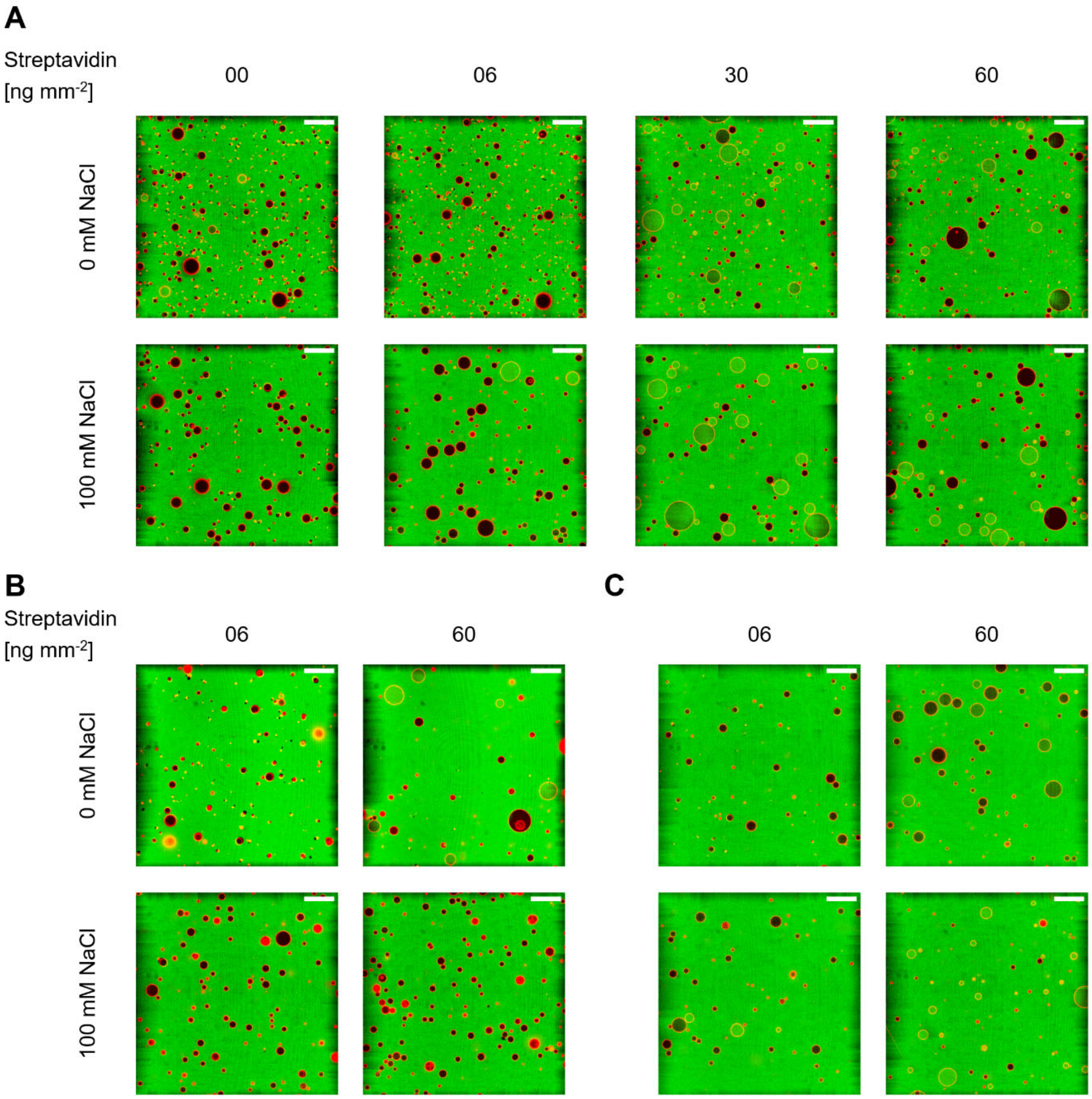


**Figure S8: Confocal microscopy images of HPTS leakage in GUVs in the presence and absence of 100 mM NaCl.** Overlay of HPTS channel (green) and Liss Rhod PE channel (red). Images were recorded approximately 5 µm above the slide surface. The scale bar is 50 µm. A) Images of immobilization-induced leakage of HPTS in PVA GUVs at different streptavidin densities. B) Images of immobilization-induced leakage of HPTS in Pt wire GUVs at low and high streptavidin density C) Images of leakage of HPTS in PVA GUVs with addition of dye after immobilization at low and high streptavidin density

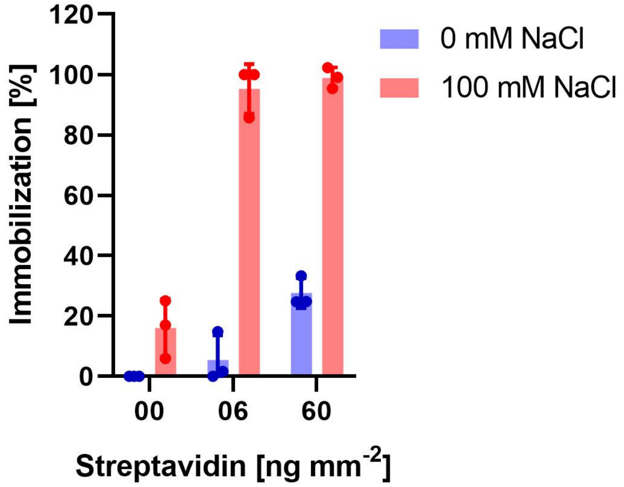


**Figure S9 Immobilization assay using channel slides and PVA GUVs with 5 mM MOPS-KOH buffer.** Percentage of immobilized PVA GUVs in the presence or absence of 100 mM NaCl at different streptavidin densities assessed by comparing the number of immobilized GUVs and the number of GUVs before application of flow. Data from three experiments are shown. GUVs were prepared in 5 mM MOPS-KOH pH 7.4, 200 mM sucrose with salt as indicated. The height of the bar indicates the average percentage of immobilization with individual values from the experiments shown as dots. Error bars indicate the standard deviation.

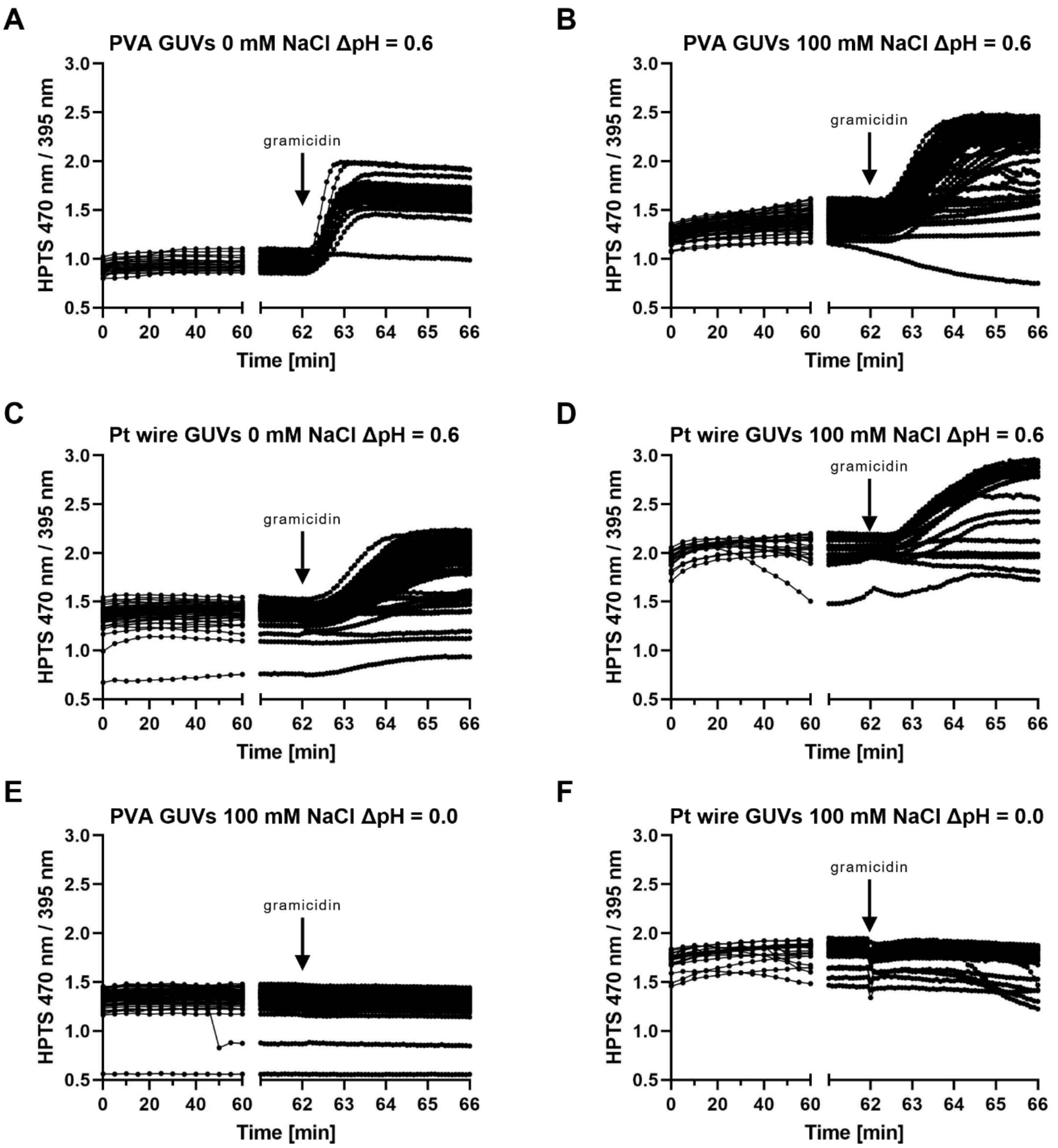


**Figure S10: Proton leakage of immobilized PVA and Pt wire GUVs in the presence or absence of 100 mM NaCl.** Analysis was limited to GUVs with diameters of 5 – 20 µm in the focal plane. Data are taken from a single time series for each condition. Each time point is represented by a dot. HPTS ratio of individual GUVs from one experiment are shown. Only GUVs with diameters of 5 – 20 µm in the focal plane were analyzed. A - D shows GUVs subjected to a pH gradient of 0.6 (pH inside = 7.4, pH outside = 8.0). E and F shows GUVs in the presence of 100 mM NaCl with no applied pH gradient. After 62 min, gramicidin was added to equilibrate the inner and outer pH. A) PVA GUVs in the absence and B) in the presence of 100 mM NaCl. C) Pt wire GUVs in the absence and D) in the presence of 100 mM NaCl. E) PVA GUVs and F) Pt wire GUVs without pH gradient.

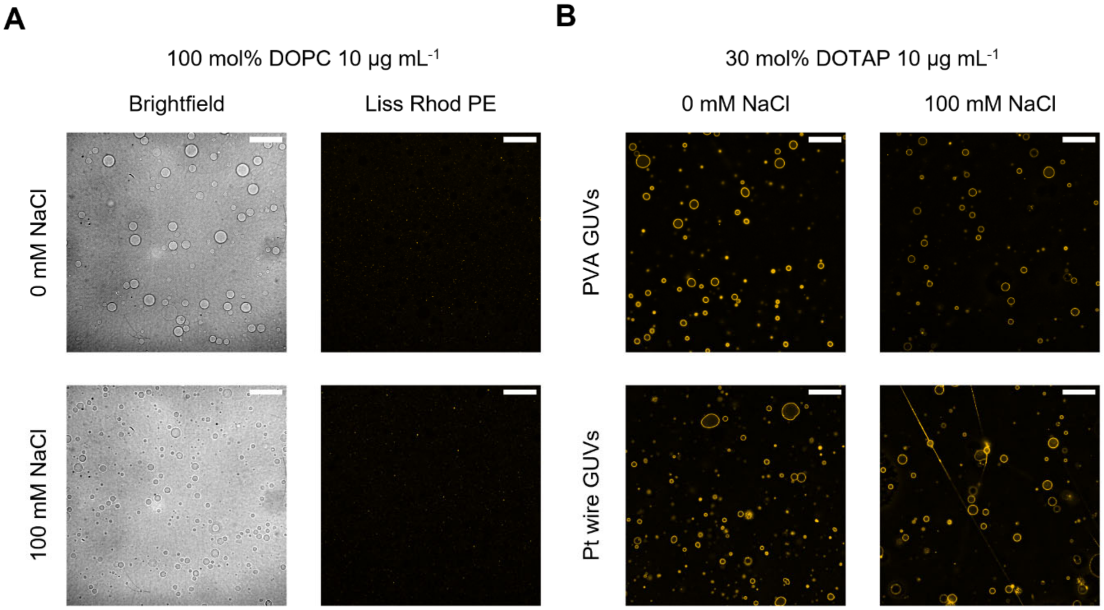


**Figure S11: Representative confocal microscopy images of charge-mediated fusion performed in an 8 well chambered slide between negatively charged GUVs and neutral or positively charged SUV.** The scale bar is 50 µm. All images from each channel were processed identically. A) Charge-mediated fusion with PVA GUVs and neutral SUVs (10 µg $mL^{-1}$) in the presence or absence of 100 mM NaCl. Brightfield and Liss Rhod PE images are shown. No fusion was observed, indicated by the absence of visible GUVs in the Liss Rhod PE channel. B) Charge-mediated fusion with PVA and Pt wire GUVs and positively charged SUVs (10 µg $mL^{-1}$) in the presence or absence of 100 mM NaCl.

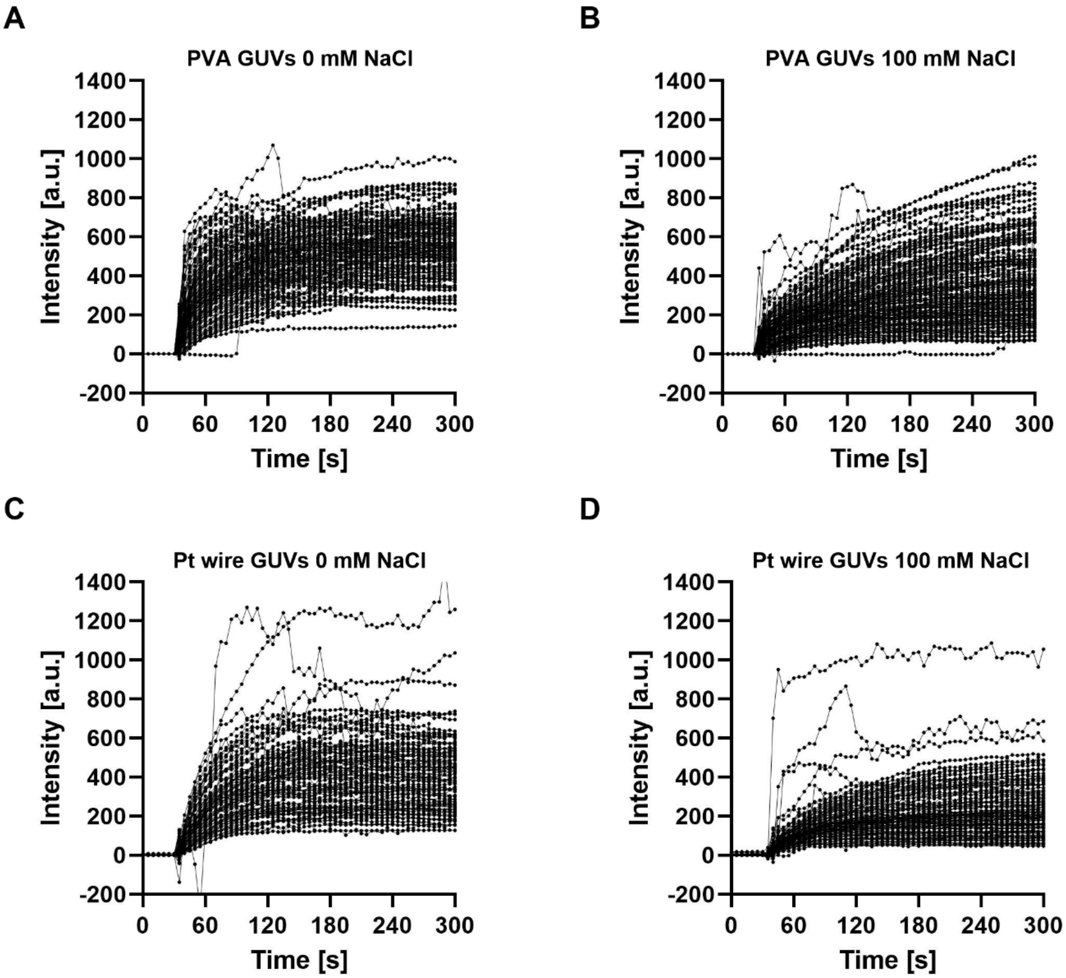


**Figure S12: Charge-mediated fusion performed in an 8 well chambered slide between negatively charged GUVs and positively charged SUVs (10 µg mL$^{-1}$).** Combined traces of GUVs from four experiments are shown. Only GUVs with diameters of 5 – 20 µm in the focal plane were analyzed (90 – 170 GUVs for all four experiments). Each recorded time point is represented as a dot. A) Fusion with PVA GUVs in the absence and B) in the presence of 100 mM NaCl. C) Fusion with Pt wire GUVs in the absence and D) in the presence of 100 mM NaCl.

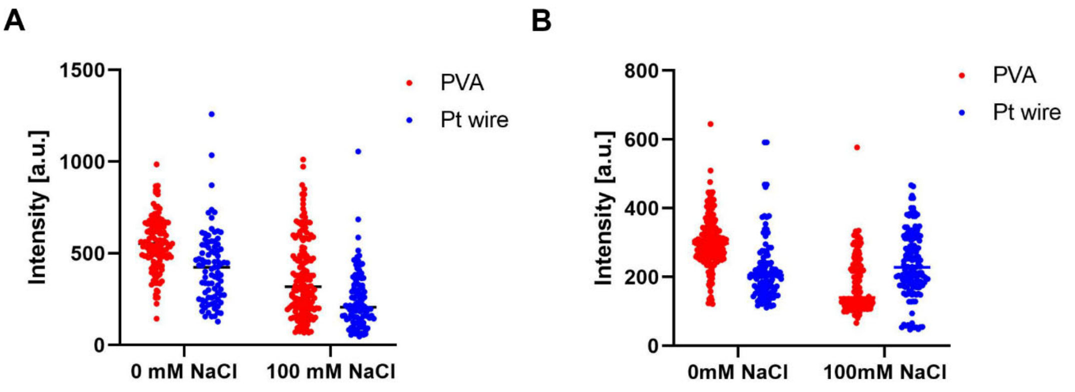


**Figure S13: Charge-mediated fusion between negatively charged GUVs and positively charged SUVs.** Combined distribution of Liss Rhod PE intensities from four experiments for PVA and Pt wire GUVs in the presence or absence of 100 mM NaCl. A) Intensity distribution from a single imaging plane after 270 s of fusion in an 8 well chambered slide. Concentration of SUVs in the slide was 10 µg $mL^{-1}$. B) Intensity distribution from an average projection of a Z-stack after fusion in an Eppendorf tube. Concentration of SUVs in the tube was 10 µg $mL^{-1}$. GUVs were fused for 15 min in the tube, added to an 8 well chambered slide coated with BSA and imaged after they were left to settle for 1h.

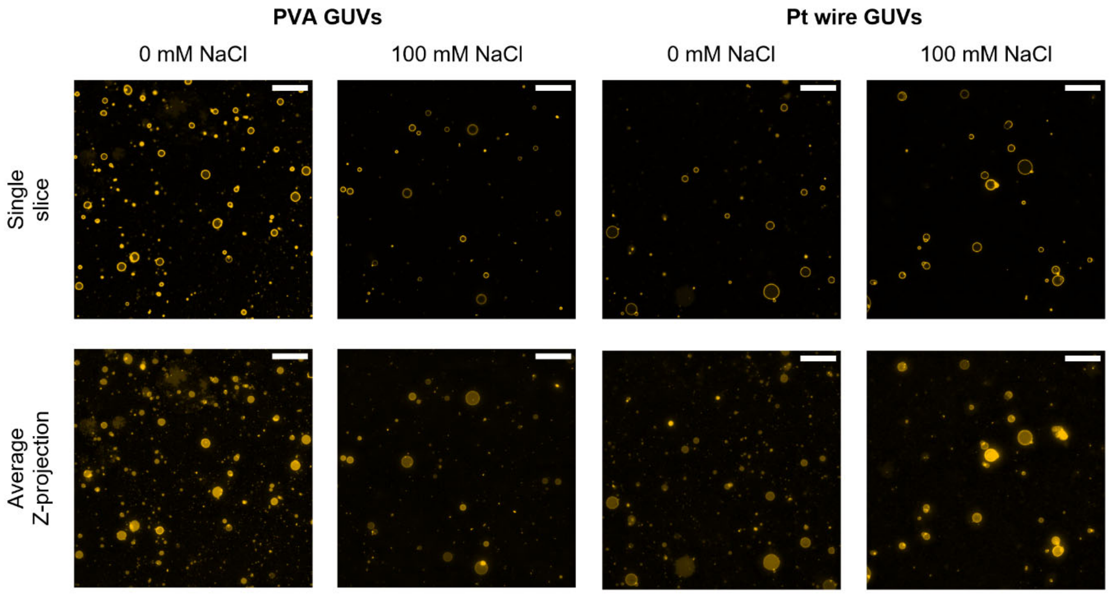


**Figure S14: Representative confocal microscopy images of charge-mediated fusion between negatively charged GUVs and positively charged SUV performed in an Eppendorf tube.** A single slice and the average Z-projection from a Z-stack image are shown for fusion with PVA and Pt wire GUVs and positively charged SUVs in the presence or absence of 100 mM NaCl. Concentration of SUVs in the tube was 10 µg $mL^{-1}$. GUVs were fused for 15 min in the tube, added to an 8 well chambered slide coated with BSA and imaged after they were left to settle for 1h. The scale bar is 50 µm. All single slice images were processed identically. All average Z-projections were processed identically.

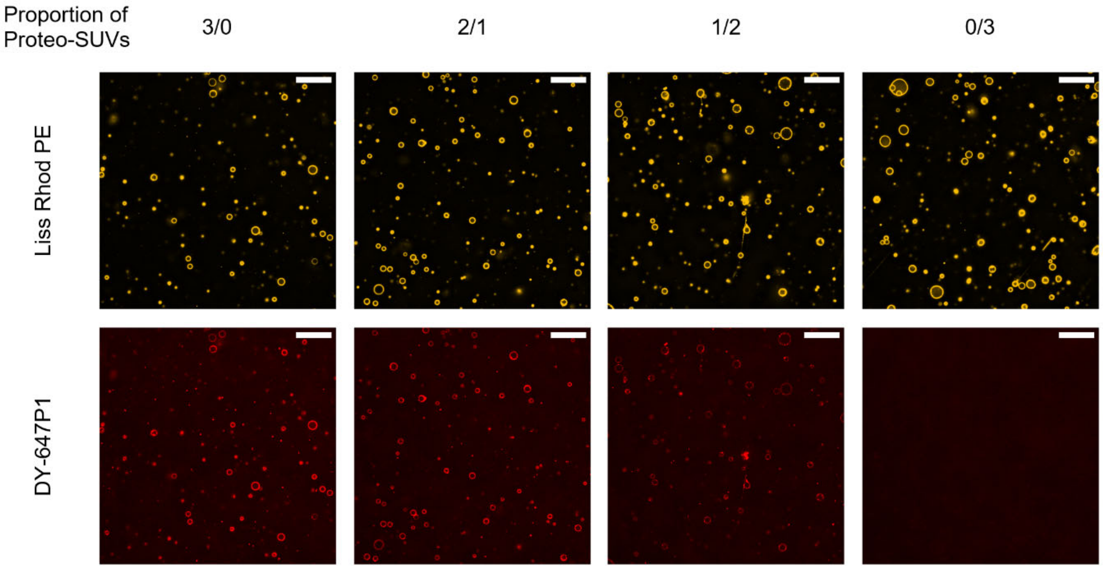


**Figure S15: Confocal microscopy images of PVA GUVs at 0 mM NaCl fused with empty and proteo-SUVs (final concentration 40 µg mL$^{-1}$) containing DY-647P1-labeled cytochrome *bo*$_3$ ubiquinol oxidase.** Empty and proteo-SUVs were mixed at different ratios with proteo-SUV proportions of 1, 2/3, 1/3 and 0. Both the Liss Rhod PE and DY-647P1 channel are depicted. The scale bar is 50 µm. All images from each channel were processed identically.

**Supplementary Movie S1:** Charge-mediated fusion of rhodamine labeled positively charged empty SUVs with negatively charged GUVs in the microscopy well using 10 mM MOPS-BTP pH 7.4, 100 mM NaCl, and 200 mM glucose. A time series was recorded for 60 s with 1 s intervals using a 40x objective (scale bar 50 µm). The movie is displayed at a rate of 5 frames per second. SUVs were added after 6 seconds.

**Supplementary File S1:** The Autodesk Fusion 360 Archive File (*.f3d) of the slide holder used for ITO electroformation.
